# A specialized cold sensing system in the naked mole-rat

**DOI:** 10.1101/2025.09.25.678629

**Authors:** Lin Wang, Alejandro Gómez-Restrepo, Amanda Niqula, Jean-Sebastien Jouhanneau, Daniel Mendez-Aranda, Wenhan Luo, James F.A. Poulet, Elvira de la Peña, Félix Viana, Valérie Bégay, Gary R. Lewin

## Abstract

Avoiding cold or seeking warmth are universal animal needs. Homeothermic mammals maintain a constant body temperature despite fluctuating ambient temperatures. One exception is the naked mole-rat which lacks effective thermogenesis. We show that the naked mole-rat has evolved a greatly expanded cold sensing system. Compared to mice, this species has many more cold-sensitive sensory neurons and most express the cold-activated TRPM8 channel. The naked mole-rat TrpM8 gene harbors a unique upstream exon that when translated produces a TRPM8 protein with a 71 amino acid N-terminal extension. When expressed, the N-terminal extension prevents membrane targeting and abolishes TRPM8 function. Splice forms lacking the extension formed functional cold and menthol activated channels *in vivo*. Additionally, many naked mole-rat sensory neurons use a TRPM8-independent mechanism to detect cold. Thus, we identified molecular changes that confer both sensitivity and flexibility to cold sensing in a species that is critically dependent on following thermal cues.

## Introduction

Sensing the thermal environment is of crucial importance for all species. However, different vertebrates are adapted to very warm, temperate or extremely cold environments. Mammals are in general homeotherms that are able to maintain a constant body temperature in the face of large changes in external temperature. However, animals need to perceive environmental temperature to make behavioral adjustments that benefit their overall homeostatic state. For example, seeking shade under extreme daytime temperatures. At another level we also know that humans and animals can feel the temperature of objects they touch with amazing precision being able to perceive temperature changes of around 0.5°C^1–3^. Thermal sensing is largely dependent on cutaneous afferent input from sensory neurons that respond to both cooling and warming stimuli^3,4^. Sensory neuron thermal detection depends on the expression of temperature sensitive transient receptor potential ion channels (TRP) at peripheral endings in the skin^3,5,6^. One of the best characterized of the thermosensitive TRPs is the cooling and menthol gated ion channel TRPM8^7,8^. Not only is TRPM8 expressed specifically in thermosensitive afferents but its presence is also required for the ability of mice to respond to cold^1,9–11^. Paradoxically, TRPM8 is also crucial for the perception of warm, because warming can silence cool evoked activity in a TRPM8 dependent manner^2^.

Interestingly, the TRPM8 channel has undergone evolutionary selection that has altered its threshold for thermal activation in a direction appropriate for species living in extreme temperature habitats. For example, the activation thresholds for TRPM8 channels from the cold tolerant emperor penguin and the hibernating ground squirrel are shifted to colder temperatures compared to those of the mouse and human channels^12,13^. In poikilothermic vertebrates like amphibians there is also evidence of altered TRPM8 thresholds that tune the channel to be more cold tolerant^14^. The lack of thermogenesis in amphibians and reptiles means that their core body temperature tracks ambient temperature. Homeothermic mammals are not normally faced with this problem. However, there is one unique rodent species, the naked mole-rat (*Heterocephalus glaber*), that lacks sufficient thermogenic capacity to maintain body temperature at a set point^15–19^. Naked mole-rats are highly social animals that live in colonies with a hierarchical structure with a single breeding queen, a system referred to as eusocial^15,20,21^. The burrow systems of the naked mole-rat in their native East African habitats are extensive and complex and are largely protected from large temperature fluctuations^22^. However, it is thought that naked mole-rats maintain a constant body temperature partly through huddling behavior^23,24^. Thus, naked mole-rats will seek warmer areas and will then deliver the warmth to other animals in the colony. We hypothesized that this interesting behavior requires a novel thermal sensory system shaped by molecular and physiological adaptations involving TRPM8 channels. Indeed, we show that, compared to mice, the naked mole-rat has an expanded thermal sensory system specialized to detect cooling. TRPM8 channels in the naked mole-rat are also partly expressed as a novel isoform with an extended N-terminal extension that functions to silence the channel. Many sensory neurons in the naked mole-rat with fast conducting myelinated axons are exquisitely sensitive to cooling of the skin, thus representing an additional cool sensing system, greatly expanded compared to mice or humans. Finally, although many naked mole-rat neurons detect cold in TRPM8-dependent manner there is a greatly expanded population of neurons that detect cooling independently of TRPM8 in this species.

## Results

### Behavioral preference for warm in naked mole-rats

To assess thermal preferences in naked mole-rats, we conducted a 12-zone temperature gradient ring assay ranging from 10 °C to 48 °C^25,26^. A cohort of male and female (*n*=16) animals was allowed to freely explore the ring for 40 minutes. Naked mole-rats spent more time in the warmer quadrants, with a population peak preference centered at 41 °C (Fig. 1A,B), indicating a pronounced bias toward elevated temperatures. This temperature preference was at least 7 °C higher than for wild type mice (*n*=16, 8 females, 8 males) tested in the same apparatus (Fig. 1B and fig S1F,G), consistent with previous studies^25,26^. Naked mole-rats can vary considerably in weight and larger animals tend to belong to higher ranks within the colony^21,27^. Since naked mole-rats show minimal thermogenesis^15,16,18,19^ we reasoned that body mass would influence how well heat is retained with smaller animals being more prone to losing heat. We thus stratified our animals by body mass and found that the lighter animals (<45g) exhibited a significantly higher preferred temperature (∼44°C) than their heavier counterparts (>45 g, n=7; ∼39°C, *p* < 0.05; Fig. 1C,D). Both heavier and lighter animals exhibited similar locomotor activity (fig S1A) This trend was confirmed in group-wise heatmaps and binned occupancy (fig. S1B). In contrast, no significant sex-based differences were observed in either peak temperature preference or exploration metrics (fig. S1 C-E). These findings suggest that naked mole-rat heat seeking behavior may be correlated with their ability to retain warmth according to body mass.

**Figure 1.**
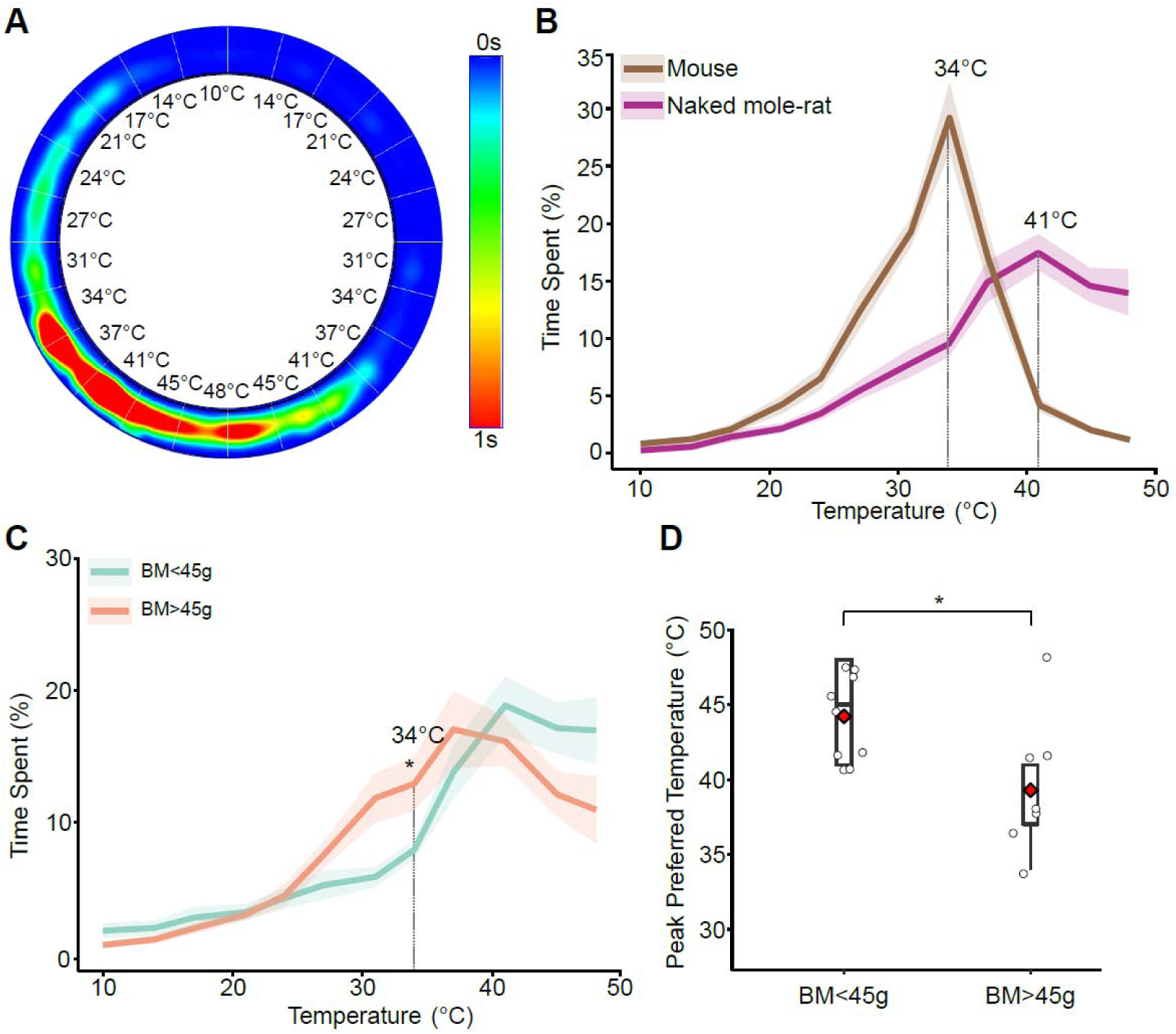
Warm-seeking behavior in naked mole-rats. (**A**) Representative occupancy heatmap of pooled data from all animals (*n* = 16; 8 males, 8 females) over 40 minutes in a 12-zone thermal gradient ring. (**B**) Group-averaged time distribution across temperature zones (mean ± SEM). Peak preference occurred at 41 °C. (**C**) Temperature occupancy curves for animals with body mass (BM) <45 g (*n* = 9) versus >45 g (*n* = 7). Asterisk indicates statistically significant group difference at 34 °C (unpaired two-tailed t-test, \**p* < 0.05). (**D**) Peak preferred temperature (per animal) grouped by body mass. Red diamonds indicate group means. Unpaired two-tailed t-test, \**p* < 0.05.

### Expanded TRPM8 expression in naked mole-rat somatosensory system

Warm-seeking behavior may also be associated with avoidance of cold temperatures which is known in many species to depend on the cooling and menthol sensitive channel TRPM8^9,12–14,28^. We next made a comparative study of the expression and distribution of TRPM8 channels in the naked mole-rat compared to the mouse. We used specific TRPM8 antibodies that were validated using dorsal root ganglia (DRG) from *Trpm8*^⁻^*^/^*^⁻^ mice^9^, in which no fluorescence signal was detected (fig. S2A). Immunofluorescence studies were further complemented with single molecule in situ fluorescence hybridization (smFISH). We used PGP9.5 as a marker of all sensory neurons within the ganglia. In mouse lumbar DRG we found that 6% of the PGP9.5-positive neurons were strongly TRPM8-positive which is in close agreement with the literature^7,28,29^. However, in the naked mole-rat lumbar DRG we found that 16% of the neurons were TRPM8-positive, significantly higher than in mouse (*p* < 0.001; Fig. 2A).

**Figure 2.**
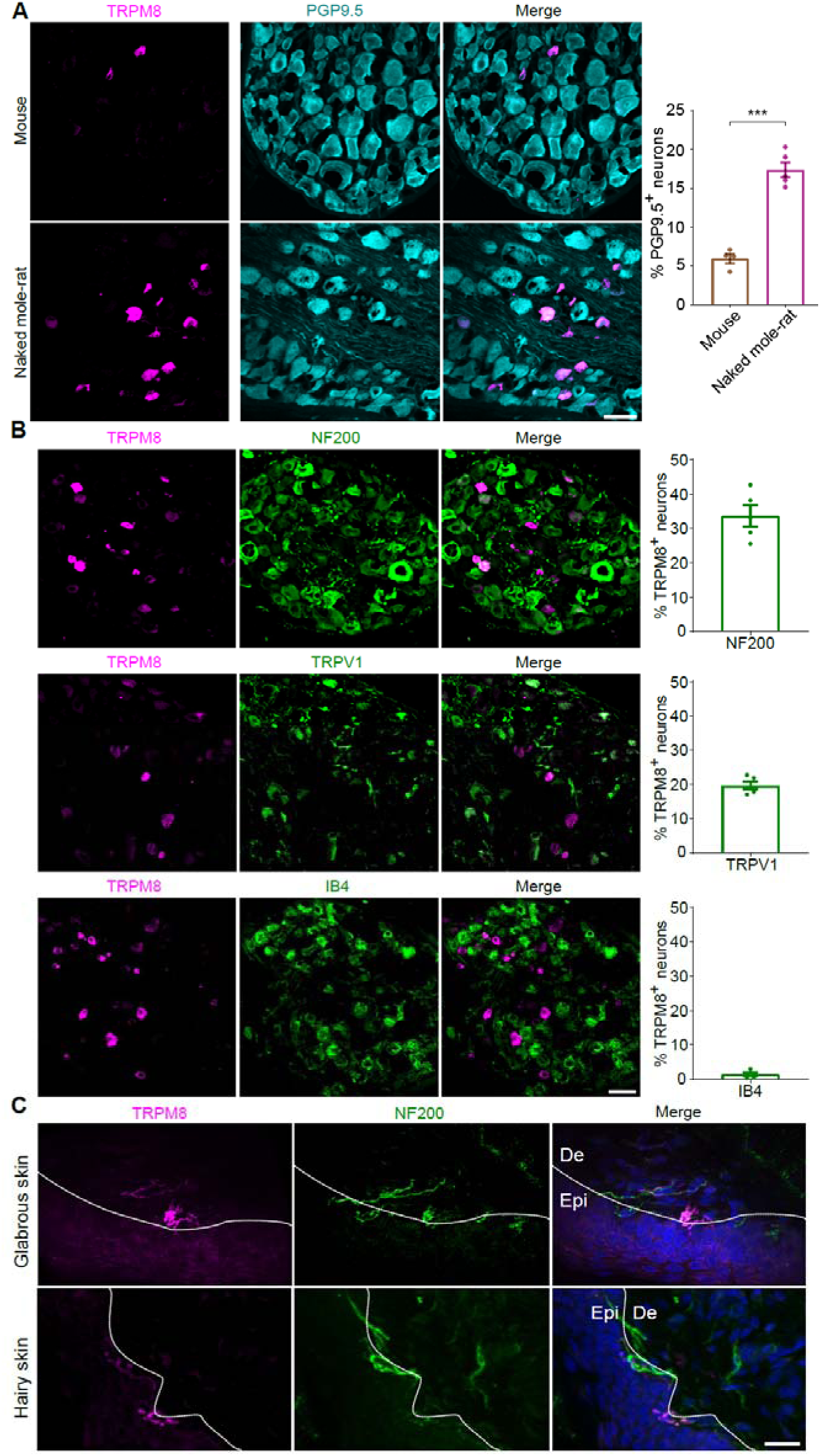
TRPM8 expression in the naked mole-rat. (**A**) Representative immunofluorescence images of dorsal root ganglia (DRGs) from adult mice (top) and naked mole-rats (bottom), stained for TRPM8 (magenta) and the pan-neuronal marker PGP9.5 (cyan). Quantification (right) shows the percentage of TRPM8⁺ neurons among PGP9.5⁺ cells. Data represent mean ± SEM. Group comparison by unpaired two-tailed Student’s t-test, \*\*\**p* < 0.001; *n* = 4 mice, *n* = 5 naked mole-rats; >4000 neurons analyzed per condition. Scale bar: 50 μm. (**B**) TRPM8⁺ neurons in naked mole-rat DRGs co-expressing molecular subtype markers. Representative images show TRPM8 (magenta) co-stained with NF200 (top, green; myelinated neurons), TRPV1 (middle, green; peptidergic nociceptors), or IB4 (bottom, green; non-peptidergic nociceptors). Quantification (right) indicates the proportion of TRPM8⁺ neurons co-labeled with each marker. *n* = 5 naked mole-rats; >2000 neurons analyzed per condition. Scale bar: 50 μm. (**C**) TRPM8 expression in naked mole-rat glabrous (top) and hairy (bottom) skin. Immunostaining shows TRPM8⁺ fibers (magenta) co-localizing with NF200⁺ fibers (green) in both the epidermis (Epi) and dermis (De). White dashed lines indicate the boundary between epidermis and dermis. Scale bar: 20 μm.

Subtype analysis revealed that TRPM8⁺ neurons were broadly distributed across sensory populations in naked mole-rats. Co-expression with NF200 (33.7%) indicated substantial expression in myelinated afferents, while partial overlap with TRPV1 (19.5%) and CGRP (39.3%) suggested localization in some polymodal nociceptors (Fig. 2B, fig. S2B). In contrast, minimal co-localization with IB4 (1.3%) showed that TRPM8 was virtually absent from non-peptidergic neurons. Note that published studies as well as our present data indicate that there is minimal expression of TRPM8 in myelinated afferents in the mouse^7,29^ (fig. S2). In the naked mole-rat skin we could also detect TRPM8-positive fibers in both epidermis and dermis, and some of these fibers also stained positive for the myelinated axon marker NF200⁺ (Fig. 2C). In the spinal cord, TRPM8 antibodies labelled the central terminals of primary afferents mostly in the superficial dorsal horn (lamina I) with some sparse labeling in deeper dorsal horn (fig. S2C–E).

### A species-specific TRPM8 isoform with an extended N-terminal domain

Examination of the naked mole-rat genome (NCBI RefSeq assembly GCF_000247695.1) revealed a novel protein coding exon at the beginning of the gene (Fig. 3A, fig S3A). Comparative sequence analysis of TRPM8 orthologs revealed that naked mole-rats express a previously undescribed variant containing a unique N-terminal extension. Using previously obtained RNAseq data from naked mole-rat DRG^30^ we found RNA reads that mapped to the novel 5í exon which lies upstream of the first protein coding exon in the mouse and human *trpm8* genes (fig. S3B). This novel exon, located upstream of the conserved melastatin homology regions (MHR1–4), was absent from mouse, human, guinea pig, and frog *Trpm8* genomic sequences (Fig. 3A, fig. S3A). Alignment of multiple amino acid sequences revealed that apart from the N-terminal extension, the core naked mole-rat sequence contained highly conserved functional domains containing all key residues required for menthol and cold activation, ligand interactions, and ion conduction (fig. S3A).

**Figure 3.**
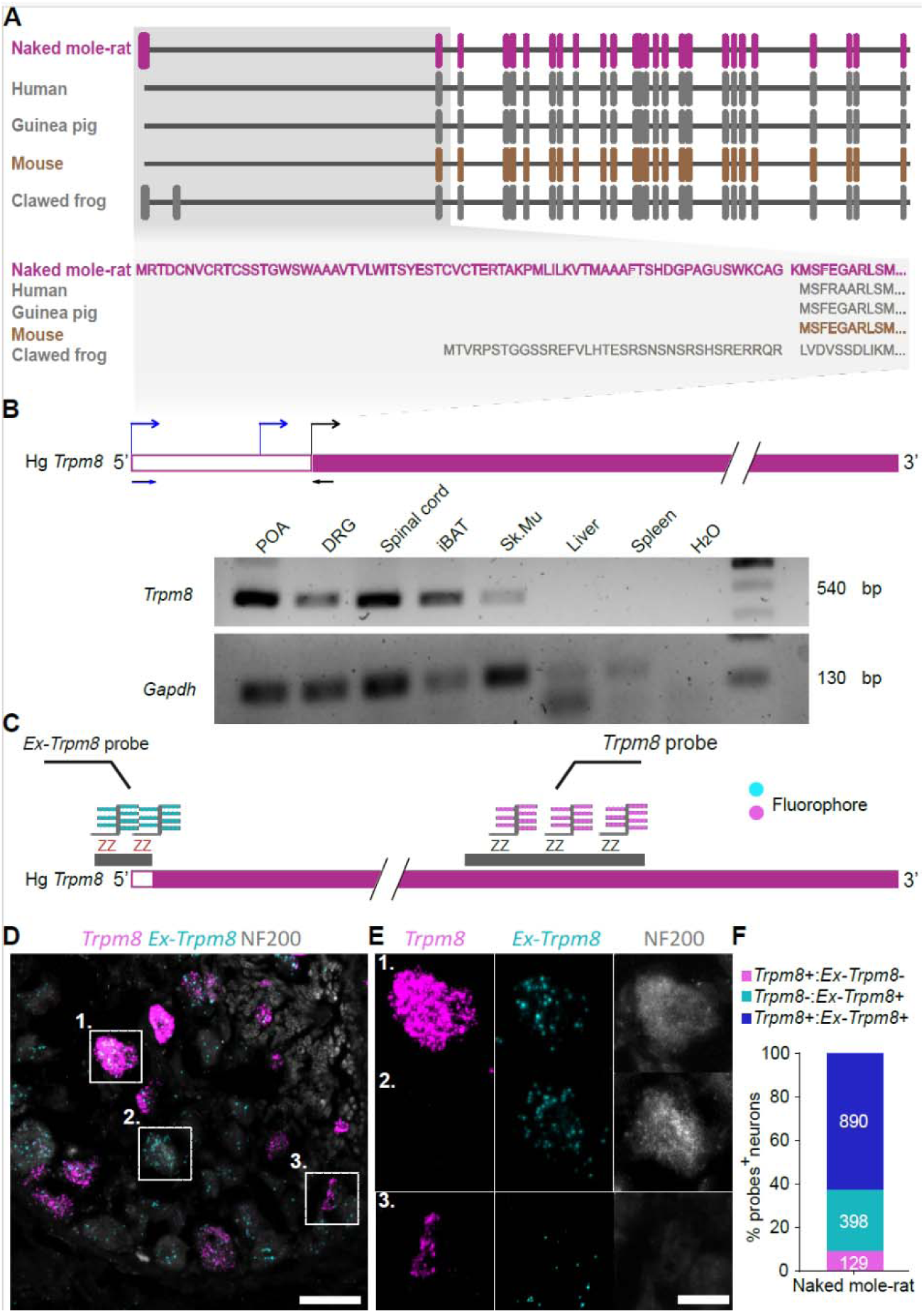
A unique TRPM8 isoform in naked mole-rat. (**A**) Comparative exon–intron structure and sequence alignment of *Trpm8* across species. Schematic alignment of *Trpm8* exon organization in naked mole-rats, humans, guinea pigs, mice, and clawed frogs. Naked mole-rats exhibit a unique N-terminal extension (magenta) not conserved in other species. Bottom: amino acid alignment highlighting the species-specific extended sequence in naked mole-rats. (**B**) Tissue-specific expression of *Trpm8* in naked mole-rats. RT-PCR detection of *Trpm8* transcripts in sensory (POA, DRG, spinal cord) and non-sensory (iBAT, Sk. Mu, liver and spleen) tissues. *Gapdh* was used as a loading control. POA, preoptic area; iBAT, interscapular brown adipose tissue; Sk.Mu, skeletal muscle.(**C**) Design of RNAScope probes targeting *Trpm8* isoforms. Schematic of probe annealing locations: canonical *Trpm8* probes (magenta) target the conserved coding region, while extended *Ex-Trpm8* probes (cyan) specifically detect the naked mole-rat–specific 5′ extension. (**D, E**) Expression patterns of canonical *Trpm8* and *Ex-Trpm8* in naked mole-rat DRG neurons. Representative RNAScope fluorescence images of DRG sections showing spatial localization of canonical *Trpm8* (magenta) and *Ex-Trpm8* (cyan) transcripts, with NF200 (gray) labeling myelinated neurons. Insets (E1–E3) display individual neurons expressing *Trpm8* only, *Ex-Trpm8* only, or both isoforms. Scale bars: 50 μm (**D**), 30 μm (**E**). (**F**) Quantification of *Trpm8* isoform expression in naked mole-rat DRGs. Stacked bar graph showing the number and proportion of neurons expressing canonical *Trpm8* only (129 neurons), *Ex-Trpm8* only (398 neurons), or both isoforms (890 neurons). Data pooled from *n* = 3 animals.

To confirm the expression of this extended isoform *in vivo*, we performed isoform-specific RT-PCR and smFISH experiments. Using primers designed to amplify RNA sequences resulting from splicing of the N-terminal exon to the canonical first exon we detected amplicons from sensory tissues (DRG and spinal cord). We could also detect expression of the novel N-terminal containing RNA in several other tissues like the Preoptic Area (POA), which is involved in temperature sensing^31,32^ as well as in non-sensory tissues like brown adipose tissue (BAT) and skeletal muscle. Other non-sensory tissues like the liver and spleen showed no detectable expression of the extended N-terminal containing *Trpm8* transcripts (Fig. 3B).

Multiplex smFISH was performed with two custom designed probes, one designed to detect mRNA transcribed from the novel N-terminal exon (called *Ex-Trpm8*) and another detecting mRNA transcribed from canonical *Trpm8* sequences (Fig 3C). We found that more than 60% of the DRG neurons expressed mRNA detected by both probes, the *Ex-Trpm8* and canonical *Trpm8* probes (*ExTrpm8+:Trpm8+*). In around 28% of the neurons mRNA was only detected by the *Ex-Trpm8* probe (*ExTrpm8+:Trpm8-*) suggesting that these cells do not express full length *Trpm8* transcripts. In the remaining neurons (∼9%) no signal was detected with the *Ex-Trpm8* probe, but the cells were positive for the canonical *Trpm8* probe (*ExTrpm8-:Trpm8+*) (Fig. 3D–F). These data indicated that not all sensory neurons express *Trpm8* transcripts that include sequences from the 5‘protein coding exon. Morphometric analysis showed that neurons expressing *Ex-Trpm8* were significantly larger in cross-sectional area compared to those expressing canonical *Trpm8* alone (*p* < 0.01, *p* < 0.001, one-way ANOVA; fig. S3C,D). Moreover, cell size distributions indicated that *Ex-Trpm8+* neurons predominantly occupied the large-diameter range, whereas *Trpm*8+: *Ex-Trpm*8-neurons were skewed toward smaller diameters (fig. S3C,D).

### The extended N-terminal domain suppresses TRPM8 function

To assess the functional impact of the extended N-terminal domain, we expressed TRPM8 isoforms in HEK293 cells and performed ratiometric calcium imaging using Fura-2. As expected, expression of the mouse TRPM8 (MmTRPM8) was associated with robust increases in intracellular calcium in response to cold (16 °C), as well as to the TRPM8 agonists, menthol (200 μM), and icilin (10 μM). Cells expressing naked mole-rat TRPM8 (HgTRPM8) exhibited highly attenuated responses to cold, menthol (200 μM), and icilin (10 μM) (Fig. 4A–B). In contrast, for mouse TRPM8, calcium influx to icilin was very significantly enhanced with simultaneous application of cold as has been observed previously^33^.The reduced calcium influx with canonical HgTRPM8 was consistent across all stimuli and were not due to differences in transfection efficiency, as the percentage of GFP⁺ cells was comparable across groups (fig. S4A). Notably, cells expressing HgTRPM8 showed minimal responses to cold alone, but exhibited significantly enhanced activation when cold was combined with icilin (Fig. 4B).

**Figure 4.**
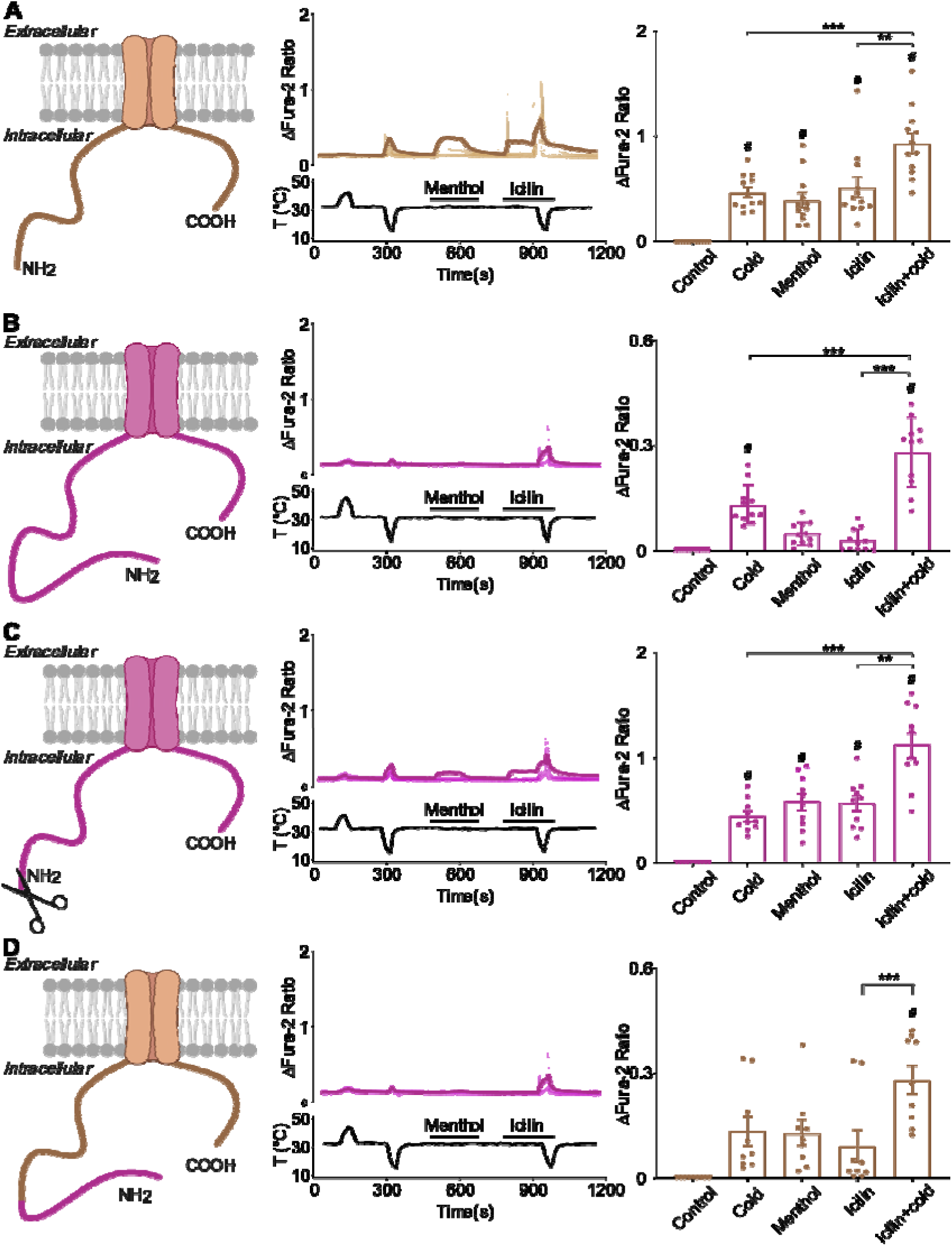
Functional analysis of TRPM8 isoforms. **(A)** Mouse TRPM8 (MmTRPM8). Top: schematic representation of mouse TRPM8 protein. Middle: representative Fura-2 calcium imaging traces showing responses to cold (18 °C), menthol (200 μM), icilin (10 μM), and combined icilin + cold stimuli. Bottom: temperature trace. Average responses in bold traces. Right: quantification of ΔFura-2 ratio under each condition. **(B)** Full-length naked mole-rat TRPM8 (HgTRPM8). Top: schematic representation including the extended N-terminal domain. Middle: representative Fura-2 traces. Bottom: temperature trace. Right: quantification of ΔFura-2 ratio. **(C)** N-terminal truncated TRPM8 (ΔN1–71 HgTRPM8). Top: schematic representation of deletion construct. Middle: representative Fura-2 traces. Bottom: temperature trace. Right: quantification of ΔFura-2 ratio. **(D)** Chimeric TRPM8 (HgNter–MmTRPM8). Top: schematic representation of chimeric construct with naked mole-rat N-terminal sequence fused to mouse TRPM8. Middle: representative Fura-2 traces. Bottom: temperature trace. Right: quantification of ΔFura-2 ratio. Quantification of ΔFura-2 ratio for each construct. Data are presented as mean ± SEM from n = 9–12 coverslips per construct, pooled from ≥4 independent transfections, with >50 transfected cells analyzed per construct. Statistical comparisons were performed using one-way ANOVA with Tukey’s post hoc test. Asterisks indicate significant differences between stimulus within each construct: \*\**p* < 0.01, \*\*\**p* < 0.001. Hash symbols indicate significant differences between stimulus and control within each construct: # *p* < 0.001. Temperature traces shown below each set of calcium recordings.

To test whether the extended N-terminal segment is responsible for the apparently reduced function, we generated a deletion construct (ΔN1–71 HgTRPM8) lacking the extended domain. Removal of the extension restored cold and agonist sensitivity to levels comparable to MmTRPM8 (Fig. 4C). Conversely, insertion of the extended N-terminal domain into MmTRPM8 (HgNter–MmTRPM8) led to a suppression of calcium signals to both cold and agonists to levels similar to those for HgTRPM8 (one-way ANOVA, *p* < 0.001; Fig. 4D). These functional changes were not explained by differences in transcription, as RNAscope analysis confirmed expression of both canonical and extended isoforms in transfected cells (fig. S4F). Note that calcium signals mediated by the naked mole-rat deletion construct (ΔN1–71 HgTRPM8) could be clearly blocked with the selective TRPM8 antagonist AMTB (N-(3-aminopropyl)-2-[(3-methylphenyl)methoxy]-N-(2-thienylmethyl)benzamide)^34^ (fig S4G). Quantification of stimulus-responsive GFP⁺ cells revealed that constructs containing the extended domain had significantly fewer responding cells compared to canonical constructs (one-way ANOVA, *p* < 0.01; Fig. S3B–E), supporting the conclusion that the extended domain suppresses TRPM8 function.

The results from our calcium imaging experiments indicated that the N-terminal extension of the naked mole-rat TRPM8 functions to strongly inhibit channel activity. However, calcium influx measurements only indirectly assess channel function and cannot give information about the voltage sensitivity or current passed by the expressed channels. We thus assessed the physiological function of the wild type and mutant channels by making whole-cell patch-clamp recordings from HEK293 cells expressing the channels (Fig. 5A-C). As expected, expression of wild type mouse TRPM8 (MmTRPM8) revealed cooling activated inward and outward currents which were greatly potentiated by concurrent stimulation with the agonists icilin and menthol (Fig. 5A-C). However, expression of full-length naked mole-rat channels (HgTRPM8) was associated with minimal or no current activation compared to mock transfected cells (Fig 5A-C). Removal of the N-terminal extension (ΔN1–71 HgTRPM8) was associated with currents with amplitudes that were almost indistinguishable from wild type MmTRPM8 channels. Finally, by adding the novel N-terminal sequences to the mouse TRPM8 channel (HgNter–MmTRPM8) we observed an almost 5-fold loss of cooling and agonist-gated currents. However, in this latter case larger amplitudes of inward and outward currents were observed concurrent agonist and cooling stimuli than for the full-length naked mole-rat TRPM8 (HgTRPM8). These data showed that addition of the N-terminal sequences to the TRPM8 abrogates all channel activity and is not specific for any one mode of activation. These data were quantified by fitting current traces and calculating the voltage at half-maximal activation (V_1/2_) (fig S5A-E), note that the full-length naked mole-rat TRPM8 (HgTRPM8) led to currents that could not be fitted.

**Figure 5.**
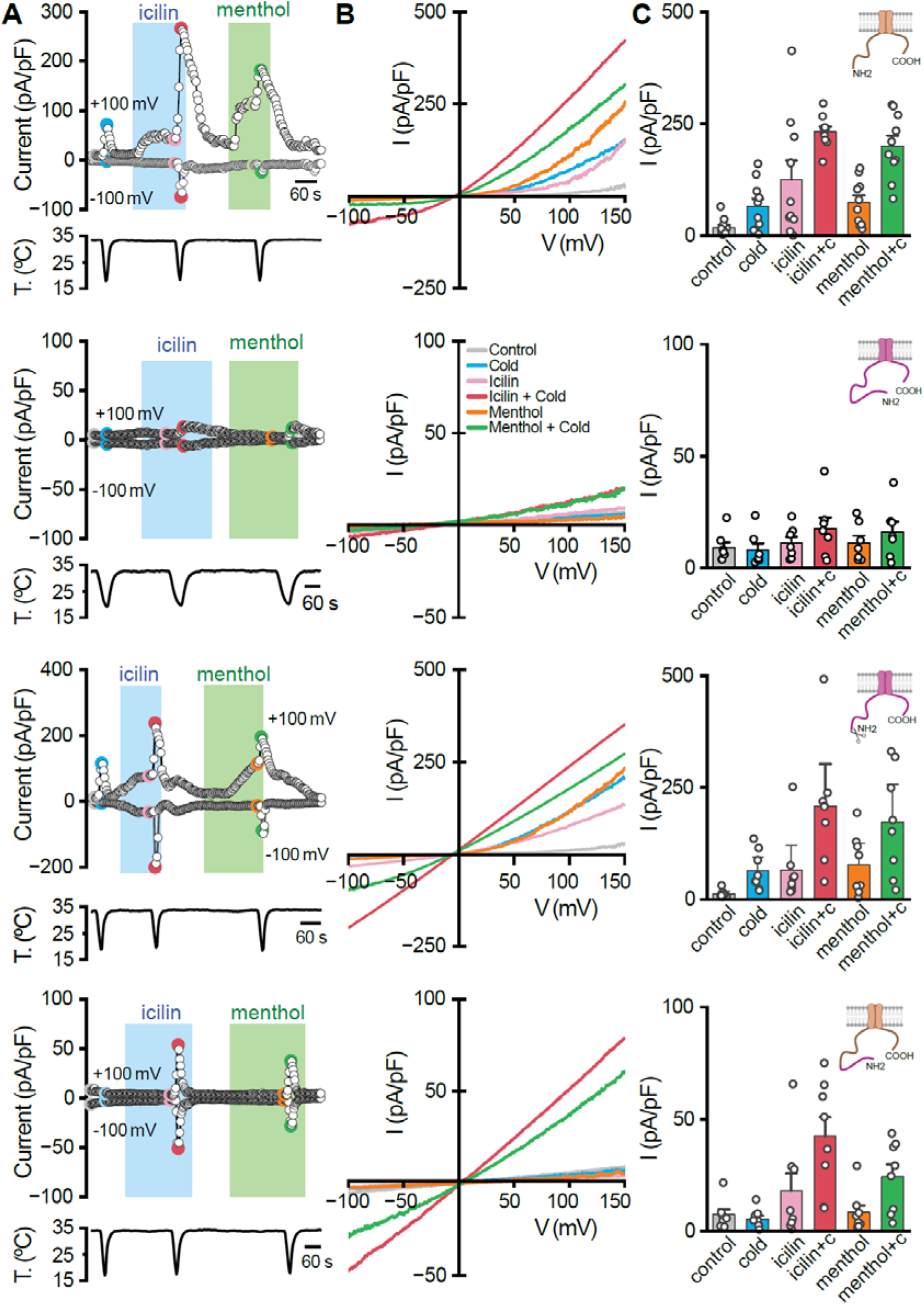
Electrophysiological validation of naked mole-rat TRPM8 hyposensitivity. (**A**) Representative time course of whole-cell currents at −100 and 100 mV in HEK293 cell transiently transfected with MmTRPM8 during application of agonists. Bottom, simultaneous recording of the bath temperature during the experiment. (**B**) I–V relationship of responses shown in **A**, obtained with a 400 ms voltage ramp from −100 to 150 mV. The color of individual traces matches the color at each particular time point in A. Note the potentiation of cold-evoked responses in the presence of icilin (10 µM) or menthol (100 µM). (**C**) Bar histogram summarizing the mean current density values at 100 and −100 mV to the different stimuli shown in **A**, with the same color code.

### The extended N-terminal domain disrupts TRPM8 membrane trafficking

Our calcium imaging experiments revealed the suppression of TRPM8 function associated with the N-terminal extension, but strong stimulation (e.g. Icilin plus cold) was still associated with significant calcium signals or membrane current in some transfected cells (Fig. 4, Fig. 5). We hypothesized that these data may be consistent with the novel N-terminus hindering the normal trafficking of channels to the membrane. We thus performed immunofluorescence imaging in HEK293T and N2a cells expressing canonical and variant constructs and used our validated TRPM8 antibody to investigate the sub-cellular distribution of TRPM8 in transfected cells. In both HEK293T and N2a cells transfected with MmTRPM8 and ΔN1–71 HgTRPM8, MmTRPM8 immunofluorescence exhibited a punctate, speckled localization pattern consistent with vesicular or membrane-associated distribution (Fig. 6A,B, fig. S6A–B). In contrast, HgTRPM8 and the chimeric HgNter–MmTRPM8 showed diffuse cytoplasmic staining, with markedly reduced speckling (*p* < 0.001; Fig.6A,B, fig S6A, B, one-way ANOVA). The TRPM8 immunofluorescence distributions were quantified using automated segmentation in FIJI, with speckled cells defined as those containing ≥2 discrete TRPM8⁺ puncta (see Methods). The speckling phenotype was restored in the ΔN1–71 construct, further implicating the extended domain in disrupting channel trafficking.

**Figure 6.**
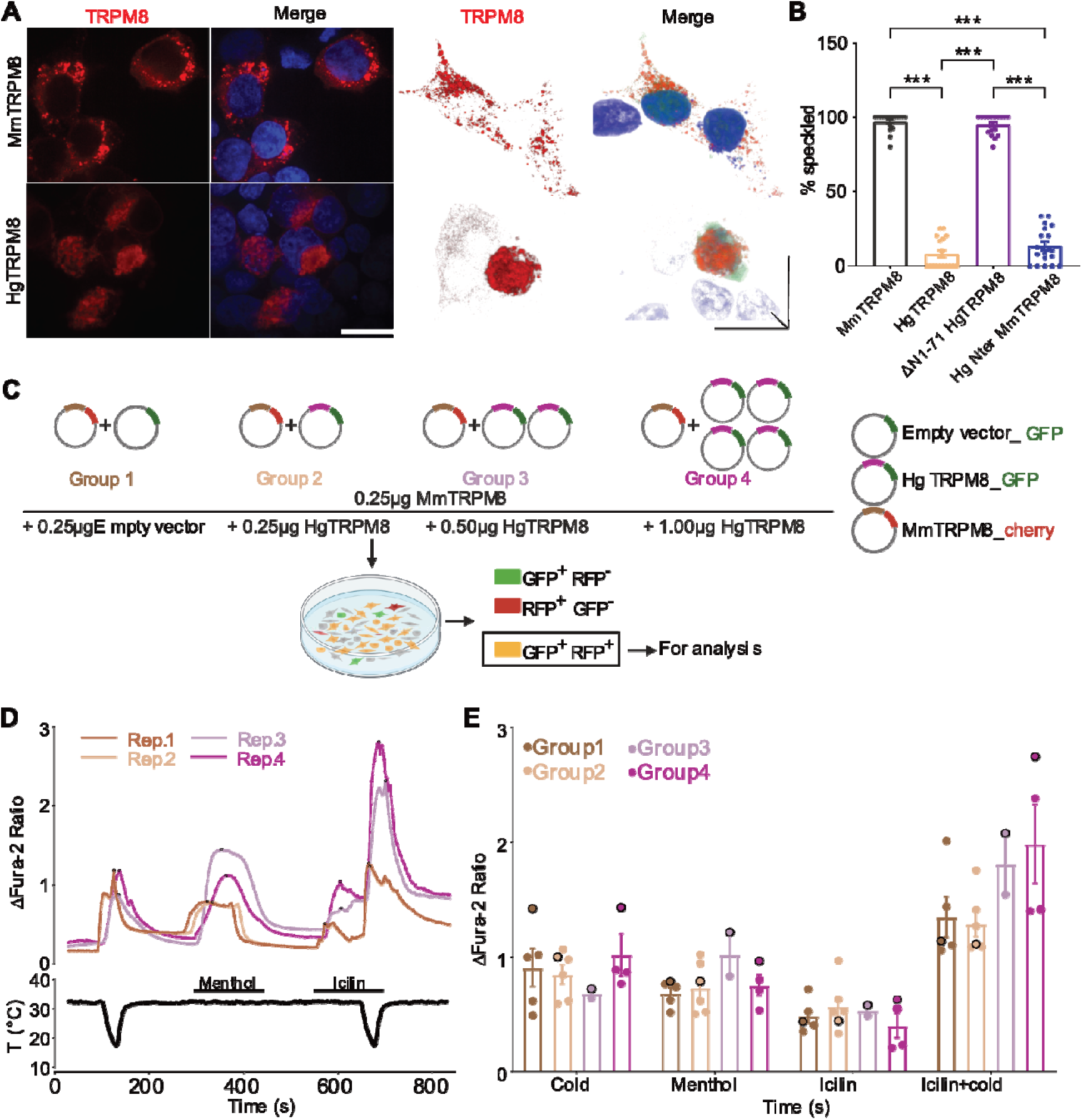
Competitive expression and functional analysis of TRPM8 isoforms. **(A)** Representative immunofluorescence images of HEK293T cells transfected with mouse TRPM8 (MmTRPM8, top) or naked mole-rat TRPM8 (HgTRPM8, bottom), stained with TRPM8 antibody (red) and DAPI (blue). Right: binary segmented images showing punctate “speckled” localization in MmTRPM8-expressing cells and diffuse cytoplasmic signal in HgTRPM8-expressing cells. Scale bar: 20 μm. **(B)** Quantification of the percentage of cells showing speckled TRPM8 staining across constructs. Data represent mean ± SEM, *n* = 30 cells per construct from ≥3 independent transfections. One-way ANOVA with Tukey’s post hoc test; \**p* < 0.001 for all indicated pairwise comparisons. Speckled cells were defined as those containing ≥2 discrete TRPM8⁺ puncta, as determined by automated segmentation in FIJI (see Methods). **(C)** Schematic of the experimental design showing four co-transfection conditions used in calcium imaging. HEK293T cells were transfected with constant amounts of mouse TRPM8 (MmTRPM8, mCherry-tagged, 0.25 μg) and increasing amounts of naked mole-rat TRPM8 (HgTRPM8, GFP-tagged; 0, 0.25, 0.5, or 1.0 μg). Cells co-expressing both GFP and RFP were selected for analysis. **(D)** Representative ΔFura-2 fluorescence ratio traces for each transfection group showing intracellular calcium responses to cold (18°C), menthol (200 μM), and icilin (10 μM). Temperature trace shown below. **(E)** Quantification of ΔFura-2 ratio in response to each stimulus across groups. Each data point represents the peak response of a single GFP⁺RFP⁺ cell; data represent mean ± SEM; *n* ≥ 2 coverslips per group from ≥2 independent transfections. One-way ANOVA with Tukey’s post hoc test; no significant differences were observed across groups under any stimulus condition (*p* > 0.05).

Three-dimensional reconstructions confirmed that speckling loss was not limited to the imaging plane (fig. S6A). This diffuse staining pattern associated with the N-terminal extension was reproduced in neuronal N2a cells, indicating that the mis-localization was not cell-type specific (Fig. S6C). These findings suggest that the extended N-terminal domain impairs the trafficking of the channel to the membrane and rather functions to retain channels in the cytosol preventing efficient plasma membrane expression. Such a function of the N-terminal extension would probably be sufficient to explain the suppression of function seen in our calcium imaging and electrophysiological experiments.

If the extended TRPM8 isoform hinders plasma membrane trafficking, it is also possible that these channels may affect the trafficking of canonical TRPM8 ion channels thus exerting a dominant negative effect. To test this idea functionally we co-expressed canonical and extended TRPM8 isoforms, maintaining the same concentration of MmTRPM8, but increasing that of HgTRPM8 to up to four times higher with transfection efficiency of each plasmid monitored by GFP or RFP expression, respectively. Functional readouts from GFP⁺/RFP⁺ double-positive cells revealed no significant differences in ΔFura-2 signals across groups, indicating that the presence of the extended isoform (HgTRPM8) does not suppress the functional expression of canonical TRPM8 channels (MmTRPM8) (Fig. 6C-E), at least in a heterologous expression system.

### TRPM8-dependent and independent cold responses

We performed calcium imaging with dissociated DRG neurons to compare cellular cold sensitivity between mice and naked mole-rats (Fig. 7A-C). The proportion of neurons responding to cold (32 °C to 16 °C) was more than three times higher in the naked mole-rat (∼20%) compared to the mouse (∼6%) (Fig. 7D). Soma size analysis revealed that cold-sensitive neurons in mice were almost all small in diameter (soma area <400 μm²), consistent with them being identical to cells that are *in vivo* cold-sensing C-fibers (Fig 7C). In contrast, naked mole-rat cold-sensitive neurons exhibited a broader size distribution and included a significantly larger fraction of medium-to large-diameter neurons (∼40% of all cold sensitive neurons with soma areas >400 μm²; *p* < 0.001) (Fig. 7C–E).

**Figure 7.**
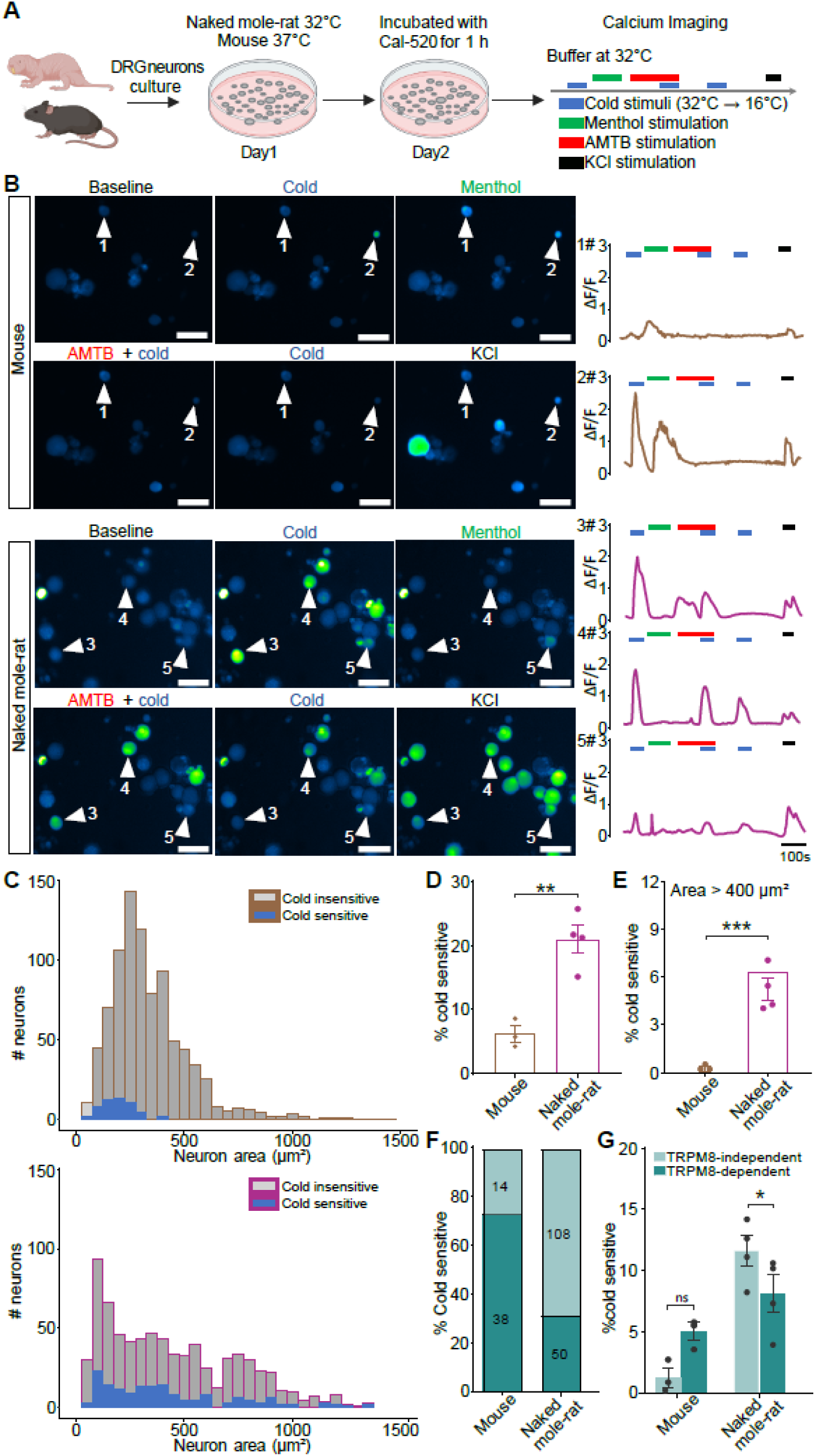
TRPM8-dependent and independent cold activation of naked mole-rat sensory neurons. **(A)** Schematic of calcium imaging workflow. Dorsal root ganglia (DRGs) from mice and naked mole-rats were dissociated and cultured overnight (mouse at 37 °C; naked mole-rat at 32 °C), then incubated with Cal-520 for 1 h before live-cell imaging. Cells were stimulated sequentially with cold buffer (32 °C → 16 °C), menthol (200 μM), AMTB (10 μM) + cold, and KCl (50 mM) at 32 °C (timeline right). **(B)** Representative fluorescence images and ΔF/F₀ calcium traces of DRG neurons from mice (top) and naked mole-rats (bottom) under baseline, cold, menthol (200 μM), AMTB (10 μM) + cold, and KCl (50 mM) conditions. Cold-responsive neurons are marked with white arrowheads and numbered. Scale bars: 50 μm. **(C)** Soma size distribution of cold-sensitive and cold-insensitive neurons in mice (top) and naked mole-rats (bottom). **(D)** Percentage of cold-sensitive neurons in mouse and naked mole-rat DRG cultures. Data represent mean ± SEM; *n* = 3 mice, *n* = 4 naked mole-rats; ≥200 neurons analyzed per animal. Group comparison by unpaired two-tailed Student’s *t*-test; \*\**P* < 0.01. **(E)** Proportion of cold-sensitive neurons with soma areas >400 μm². Unpaired two-tailed Student’s *t*-test; \*\*\**P* < 0.001. **(F)** Classification of cold-sensitive neurons as TRPM8-dependent or TRPM8-independent based on responsiveness to menthol (200 μM) and block by AMTB (10 μM). Left: stacked bar chart showing the number of cold-sensitive neurons in each category. Right: percentage of TRPM8-dependent and –independent neurons per species. **(G)** Same data as in (F), presented with individual biological replicates. Data shown as mean ± SEM; n = 3 mice, n = 4 naked mole-rats. Group comparisons were performed using unpaired two-tailed Student’s *t*-test; \**p* < 0.05, \*\**p* < 0.01, ns = not significant.

To assess TRPM8 involvement, we used the TRPM8 antagonist AMTB in DRG cultures from mice. AMTB (10µM) eliminated nearly all cold-evoked responses (∼80%), confirming TRPM8 dependence (Fig. 7F, fig S7). In naked mole-rats, however, 108 of 158 cold-sensitive neurons (∼68%) remained cold-responsive in the presence of AMTB, suggesting the existence of a substantial TRPM8-independent population. These neurons also tended to be larger in size than TRPM8-dependent cold-sensitive neurons in the naked mole-rat (fig. S7B).

To validate the specificity of these responses, we applied an alternative TRPM8 antagonist, PBMC (1-Phenylethyl-(2-aminoethyl)[4-(benzyloxy)-3-methoxybenzyl]carbamate), and observed similar partitioning into TRPM8-dependent and –independent populations (fig. S7D–E). Notably, TRPM8-independent cold responses were reproducible across two experimental protocols (acute vs. pre-incubation with antagonists), suggesting that they are not artifacts of treatment timing (fig. 7F-G and fig S7D-F).

In addition, cold-sensitive neurons unresponsive to menthol were more abundant in naked mole-rats than in mice (Fig. S7C), further supporting an enrichment of non-TRPM8-dependent cold transducers in naked mole-rats. These findings collectively indicate that cold sensing in naked mole-rats engages both TRPM8 and alternate pathways, including a population of large-diameter sensory neurons not typically associated with cold detection in other rodents.

### TRPM8-independent cold sensitivity in afferent fibers

Cell body responses to cold is a proxy for the *in vivo* ability of sensory fibers to detect changes in skin temperature^1,35^. We thus decided to make direct recordings from cutaneous cold sensitive sensory fibers in both mouse and naked mole-rat using an *ex vivo* skin nerve preparation^1,2^ (Fig. 8A). Cold-sensitive fibers were identified by their increased firing rates during a cooling stimulus applied to their receptive field. Skin temperature was measured at the unit’s receptive field and dropped to ∼10°C during (from 32 °C) application of cold ringer solution. Consistent with our calcium imaging data we found a higher proportion of cold-sensitive fibers in naked mole-rats compared to mice (Fig. 8B), and these cold-sensitive fibers included a substantial fraction of Aδ– and Aβ-fiber units (conduction velocity CV > 1.5 m/s) (Fig. 8C). In contrast, the vast majority of cold-sensitive fibers in mice had CVs in the C-fiber range (<1.5 m/s) (Fig. 8C–D; fig. S8A–C) a finding supported by an extensive literature^1,2,9,36^. We measured the instantaneous firing rates (IFRs) of single fibers during the course of the temperature change and found that peak IFRs occurred at similar temperature ranges in both mouse and naked mole-rat (Fig. 8E,F). Heatmap analyses showed that naked mole-rat cold-sensitive units spanned a wide CV range and displayed diverse temporal firing patterns (fig. S8D), consistent with functional heterogeneity. Indeed, in every case of neurons responding to cold stimuli we were able to find a mechanosensitive receptive field for the same unit. In the case of faster conducting fibers all the neurons were clearly mechanoreceptors with low mechanical thresholds (fig S7A), however, a detailed investigation of their receptor properties was not made here.

**Figure 8.**
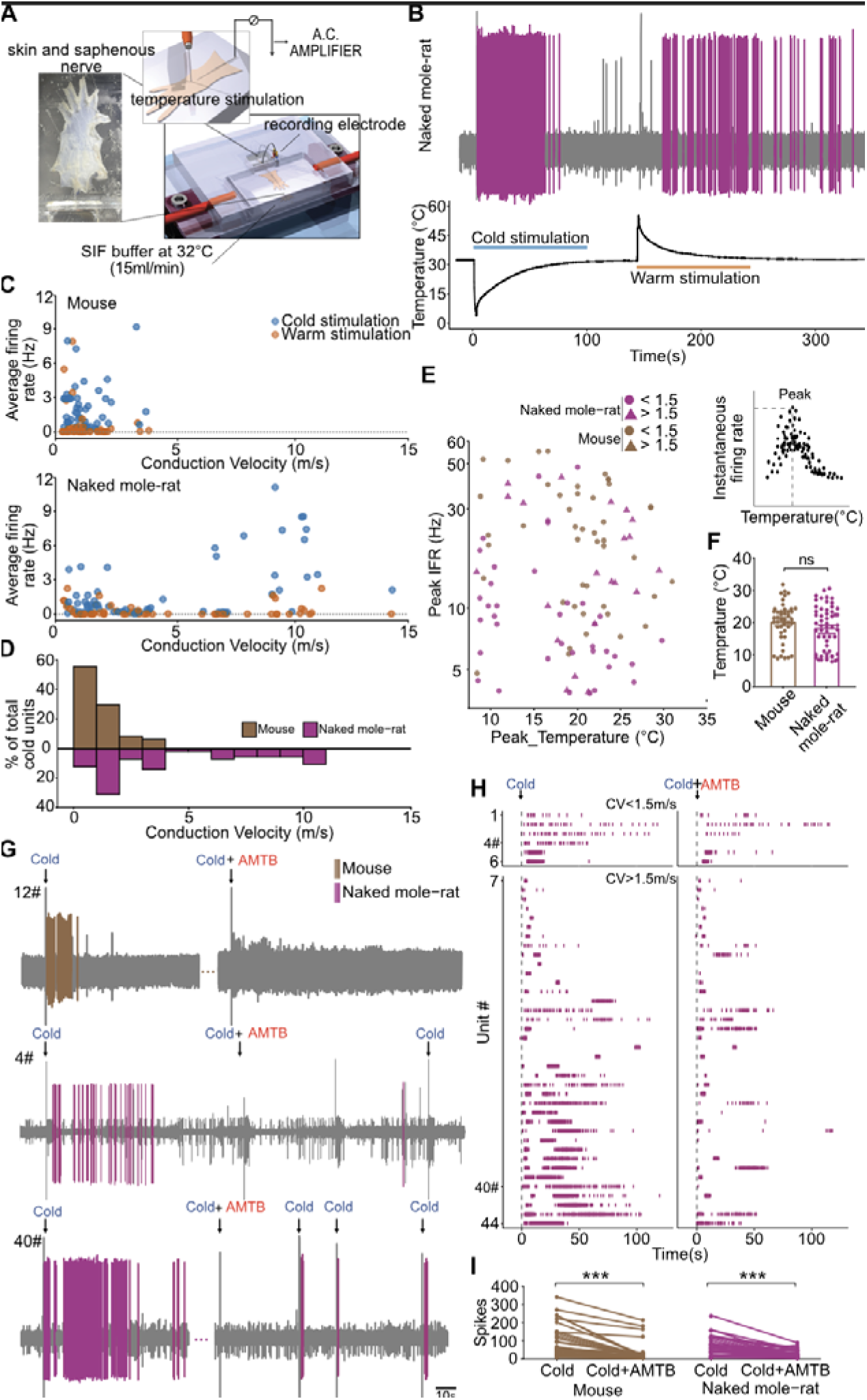
Cold activation of cutaneous sensory fibers in naked mole-rat. **(A)** Schematic of the *ex vivo* skin–nerve preparation. **(B)** Representative extracellular recording trace from a cold-sensitive fiber in naked mole-rat skin during cooling. Top: raw voltage trace of the unit. Bottom: temperature profile recorded at the receptive field. **(C)** Spike counts evoked by cold (blue) and warm (orange) stimuli across individual cold-sensitive units from mice (*n* = 47 units from 13 animals) and naked mole-rats (*n* = 59 units from 11 animals), plotted against conduction velocity (CV). **(D)** Histogram showing the distribution of conduction velocities among cold-sensitive fibers from mice (*n* = 47 units from 13 animals) and naked mole-rats (*n* = 59 units from 11 animals). **(E)** Bubble plot of peak instantaneous firing rate (IFR) versus the corresponding peak response temperature. Symbol color indicates species; symbol shape indicates conduction velocity (circle = CV < 1.5 m/s, triangle = CV > 1.5 m/s). Data represent 47 fibers from 13 mice and 59 fibers from 11 naked mole-rats. **(F)** Box plots comparing peak response temperature between mice (*n* = 47 from 13 animals) and naked mole-rats (*n* = 59 from 11 animals). Data shown as mean ± SEM. Unpaired two-tailed Student’s *t*-test; ns, not significant. **(G)** Representative firing patterns of cold-sensitive fibers from mice and naked mole-rats in response to repeated cold stimulation and AMTB (10 μM) application. Traces correspond to Unit 12 (mouse), and Units 4 and 40 (naked mole-rat), as shown in detail in panel (H) and Fig S7A–B. **(H)** Raster plots of raw voltage trace from individual cold-sensitive fibers recorded in naked mole-rats before and after AMTB application. Units are grouped by CV. Traces shown in panel (G) correspond to Unit 4 and Unit 40. A total of 44 cold-sensitive units were recorded from 8 naked mole-rats. **(I)** Paired comparison of cold-evoked spike counts before and after AMTB application in individual cold-sensitive fibers from mice (*n* = 43 units from 9 animals) and naked mole-rats (*n* = 44 units from 8 animals). Each line represents a single unit. Data shown as mean ± SEM. Paired two-tailed Student’s t-test; \*\*\**p* < 0.001.

To assess the contribution of TRPM8 to peripheral cold responses, we applied the selective antagonist AMTB onto the skin during recordings made from single cold-sensitive afferents. In mice, AMTB almost completely abolished cold-evoked firing in all recorded units (Fig. 8G–H; fig. S8A–B), consistent with a TRPM8-dependent mechanism. In contrast, a subset of cold-sensitive fibers in naked mole-rats retained robust spiking following AMTB application (Fig. 8I; fig. S8A–B), indicating the existence of TRPM8-independent cold-sensitive afferents.

## Discussion

Evolution has selected for a unique metabolic and thermoregulatory phenotype in the eusocial subterranean dwelling naked mole-rat^15,37,38^. With the possible exception of sloths (*Bradypus*)^39^, the naked mole-rat is the only known mammal that has almost completely dispensed with the ability to generate heat to maintain a constant core body temperature^16,19,40^. Naked mole-rats have a preferred body temperature of around 32°C which they maintain behaviorally e.g. by seeking warmer areas in their underground burrows^23,24^. These highly social animals also show huddling behavior that facilitates warmth retention^23,40^. We hypothesized that these unique metabolic and thermal challenges may have led to adaptations in the somatosensory system that facilitates their ability to seek warmth. Here we confirm that in comparison to mice, naked mole-rats seek warmer areas in a thermal gradient ring test (∼41 °C compared to 34 °C for mice) (Fig. 1)^40^. This behavior appears to be related to thermoregulation as smaller animals with a larger surface-to-weight ratio tend to seek even warmer areas in the ring compared to larger animals (Fig. 1C). Presumably, smaller animals seek warmer areas as they are more prone to heat loss due to their increased surface to weight ratio. Here we show that the naked mole-rat somatosensory system shows unique physiological and molecular adaptations that likely enhance the animal’s ability to avoid cold or seek warm. In particular, we show that the naked mole-rat has a considerably expanded repertoire of cold sensing sensory neurons (Fig. 2, Fig. 7, Fig. 8). This expansion includes an increase in the proportion of unmyelinated C-fiber afferents that are cold sensitive and also express the menthol activated ion channel TRPM8 (fig S8D), compared to mice (Fig. 2, Fig. 7, Fig. 8). However, we also discovered a novel naked mole-rat specific *Trpm8* isoform that is expressed by many sensory afferents in the DRG (Fig. 3,4,5). This isoform includes an N-terminal extension of the protein that is absent in closely related species and in all other animals with a *Trpm8* gene. Using calcium imaging and electrophysiology we show that the full-length naked mole-rat TRPM8 is almost completely non-functional (Fig. 4, 5). The N-terminal extension coded for by a unique 5’ exon was also shown to be necessary and sufficient to abolish channel function by preventing plasma membrane trafficking of the TRPM8 channel (Fig. 6). Finally, we have also shown that many naked mole-rat sensory afferents are very sensitive to cooling using a transduction mechanism independent of TRPM8. Together our data demonstrate how multiple molecular adaptations have allowed naked mole-rats to become exquisitely sensitive to small temperature fluctuations in their environment a trait that is probably required to compensate for their lack of thermogenic capacity.

### Expanded cold sensing system in the naked mole-rat

The naked mole-rat has been shown to be hypo-sensitive to several modalities of noxious stimuli, e.g. irritants like capsaicin, histamine and low pH^30,41–44^. Indeed, this species was also described as being behaviorally less sensitive to the TRPM8 agonist menthol^45^. One major factor that may contribute to such behaviors is the fact that the naked mole-rat is unique in having at least half the number of unmyelinated C-fibers in cutaneous sensory nerves compared to other mammals^46–48^. Here we show that this paucity of C-fibers is also reflected in the cell size distribution of acutely cultured naked mole-rat DRG neurons with many fewer small cells (<400 µm^2^ in area) as a proportion of the total compared to the mouse (Fig. 7). However, this reduction in the overall number of small diameter sensory neurons was accompanied by an overall increase in the proportion of small sensory neurons that were either cold sensitive or TRPM8-positive (Fig. 2 and Fig. 7). We have proposed that naked mole-rats have fewer nociceptors because of reduced nerve growth factor (NGF) signaling during peripheral nervous system development^46,47^. It is known that NGF can control the expression specific ion channels during sensory neuron development^49,50^. However, our data suggest that in the face of hypofunctional NGF signaling the development of the TRPM8 population is preserved, consistent with a low overlap between TRPM8 expression and TrkA in early development^51^. There is a substantial loss of small diameter sensory neurons during post-natal development in the naked mole-rat^47^, and presumably this neuronal loss spares the TRPM8 population. Furthermore, studies on the development of TRPM8 neurons in the mouse are consistent with the idea that this population is independent of NGF signaling during development^51,52^. However, we also observed a second more surprising expansion of cooling sensitive afferents with axonal conduction velocities in the Aδ range and Aβ-fiber range. Consistent with previous reports, cooling sensitive fibers with myelinated axons were very rare in the mouse^1^ (Fig. 7-8, fig S4). This unusual group of fast conducting cooling sensitive fibers in the naked mole-rat made up at least half of the total population and appeared to code cooling in a similar way to slower conducting fibers in both the mouse and naked mole-rat (Fig. 8). Interestingly, all the cooling sensitive afferents with faster conducting axons had mechanosensitive receptive fields. Although we did not investigate their receptor properties in detail, all the cold sensitive myelinated fibers appeared to be low threshold mechanoreceptors. We also routinely recorded responses of skin afferents to warming stimuli and found that both in the mouse and naked mole-rat these neurons were much less prevalent than cooling responsive neurons^2^. We observed many more TRPM8^+^ neurons in the naked mole-rat compared to mice (17% compared to 6% of all DRG neurons). However, by using TRPM8 antagonists we found that many sensory neurons in the naked mole-rat appeared to transduce cold stimuli independently of TRPM8 (∼40%) (Fig. 7-8). In contrast, the same TRPM8 antagonists almost completely abolished cool activation of mouse sensory neurons (Fig. 7, Fig. 8).

### Alternative exon usage shuts down naked mole-rat TRPM8 function

Ion channel function can be finely tuned through the expression of different isoforms that can be controlled by alternative splicing^53^. Different isoforms arising from alternative splicing have been described for several TRP ion channels, including TRPM8 channels^54–57^. Here we found that the naked mole-rat *Trpm8* gene has a unique 5’ exon coding for an N-terminal extension of the protein (Fig. 3). Using smFISH and RNA sequencing datasets we could show that transcripts encoding this novel full length TRPM8 protein are present in many DRG neurons. Functional analysis showed that in HEK293 cells the N-terminal extensions functions to almost completely abolish activation of the channel. Mechanistically, it appears that the N-terminal extension serves to prevent proper trafficking of the protein to the plasma membrane thus explaining why there appears to be an almost complete loss of TRPM8 channel function. We found evidence that some neurons express only isoforms lacking the N-terminal extension, which would lead to the expression of channels functionally indistinguishable from the mouse full length protein (fig S4). Indeed, if all transcripts coded for the full-length protein, we would not have expected to be able to record menthol activation of naked mole-rat sensory neurons in culture (Fig. 7). Co-expression of the full-length naked mole-rat TRPM8 together with mouse TRPM8 did not lead to a loss of channel function, indicating no dominant negative effects of this novel isoform. We conclude that the naked mole-rat sensory neurons may be essentially insensitive to agonists like menthol or icilin if they only express the full length TRPM8 isoform. However, after pharmacological block of TRPM8 channels we find that many sensory neurons can still transduce cold, both in culture and in the intact skin (Fig. 6). The same antagonists could be shown to effectively block naked mole-rat channels that lack the N-terminal extension (fig S4). The question naturally arises what is the TRPM8-independent transduction mechanism used by many naked mole-rat sensory neurons? There is consensus that TRPM8 is the most important cold transducing channel in mammals^3,58^, but other molecular sensors for cold have been proposed^36,59,60^. However, at the moment there is no clear consensus about what the molecular nature of TRPM8-independent cold transducers are in rodents.

We show that the addition of an upstream *Trpm8* exon in the naked mole-rat can act as a switch to effectively turn-off TRPM8 function. What could be the adaptative function of such a mechanism? Interestingly, it has been known for some time that one of the most temperature sensitive molecular mechanisms in both animal and plant cells is alternative splicing^61,62^. The naked mole-rat is unique in that body temperature fluctuates with changes in environmental temperature, and this may be the key factor leading to enhanced cold sensing in this species. Thus, a drop in temperature could in principle lead to changes in the *Trpm8* transcripts favoring functional isoforms to facilitate cold avoidance. In conclusion, we provide physiological and molecular data that reveal a uniquely enhanced cold sensing system in the naked mole-rat. Thermal information is processed in several brain areas including a thermosensory cortex in mice^63^. Our data suggest that afferent information about warm and cold is considerably expanded in this species and may be processed differently in the brain compared to homeotherms. Our finding that many mechanoreceptors are highly sensitive to cooling suggests a separate system conveying thermal information in the naked mole-rat. The function of this fast thermal system is unclear, but it may complement a more conventional C-fiber system in these animals. Naked mole-rats inhabit very complex burrow systems in the wild and need to navigate in complete darkness^64,65^. We speculate that the expanded thermosensory system of this species could also aid in navigation by enabling the animal to sense thermal gradients within their complex burrow systems.

## Materials and Methods

### Animals

Adult naked mole-rats (*Heterocephalus glaber*, 2–10 years, both sexes) and C57BL/6N wild-type or Trpm8^−/−^ mice^9^ (2–8 months, both sexes) were used in this study. Naked mole-rats were housed by colony in a system of custom-designed interconnected plastic chambers (Fräntzel Kunststoffe, Rangsdorf, Germany) within temperature– and humidity-controlled incubators (28–30 °C, 50–60% humidity) under low ambient illumination. Animals were provided ad libitum access to food; their diet consisted primarily of sweet potatoes, carrots, and celery root, supplemented weekly with ProNutro (Bokomo). Nesting material and environmental enrichment were included in all chambers. Mice were housed in individually ventilated cages (IVCs) in groups of up to five per cage under standard laboratory conditions (12 h light/dark cycle, 21–23 °C), with ad libitum access to food, water, and environmental enrichment. All experimental procedures and housing conditions were approved by the Berlin Animal Ethics Committee (Landesamt für Gesundheit und Soziales, Berlin, License: Landesamt für Gesundheit und Soziales, Berlin) and conducted in accordance with the European Union Directive 2010/63/EU on the protection of animals used for scientific purposes.

### Thermal Gradient Ring Assay

To assess thermotactic behavior in naked mole-rats, we employed a temperature gradient ring (TGR) apparatus (Ugo Basile) ^25^ modified to generate a continuous annular gradient ranging from 10 °C to 48 °C, evenly segmented into 12 discrete zones (i.e., ∼3.5 °C increment per segment). Naked mole-rats were allowed to freely explore the arena for 40 minutes during the light phase under dim illumination. A total of 16 adult naked mole-rats (N = 8 males, N = 8 females) were tested individually, and each animal’s body mass (BM) was recorded prior to the assay.

Animal movement and position were recorded using an overhead infrared camera system, and locomotor parameters—including average speed, number of line crossings, total distance traveled, and total trial entries—were quantified using custom tracking software. Time spent in each temperature zone was calculated and normalized to the total assay duration to assess thermal preference.

Thermal preference was assessed by two approaches: (1) by sex (male vs. female), and (2) by body mass. For body mass comparisons, animals were stratified into two groups: BM < 45 g (n = 9) and BM > 45 g (n = 7). Time-occupancy curves were generated for each group, and the peak preferred temperature was determined per animal as the temperature bin with the maximum occupancy.

### RNA extraction and real-Time PCR

Total RNA was extracted from dorsal root ganglia (DRG), spinal cord, preoptic area (POA), skeletal muscle, interscapular brown adipose tissue (iBAT), liver, and spleen of adult naked mole-rats using the ReliaPrep™ RNA tissue Miniprep System (Promega), according to the manufacturer’s protocol. RNA integrity was assessed using an Agilent bioanalyzer system (Agilent Technologies). Total RNA was reverse transcribed at 42 °C by using the GoScript™ Reverse Transcriptase Kit (Promega) according to the manufacturer’s protocol. Real-time PCR was run on a BioRad cycler. The following primers were used to amplify the naked mole-rat *Trpm8* specific N-terminal sequence (forward primer: 5’-CTT TGG ATC ACA TCC TAT GAA-3’ and reverse primer 5’-TCA AAC TGA ATG TCC CCG AA-3’ and Gapdh (forward primer 5’-CAA AAG GGT CAT CAT CTC TGC-3’ and reverse primer 5’-AGC AGT TGG TGG TGC AGC TA-3’). Gapdh was used as a housekeeping gene for loading control.

### Tissue collection and processing

Mice were euthanized by cervical dislocation, and naked mole-rats were sacrificed via decapitation. Dorsal root ganglia (L3–L5), lumbar spinal cord, and glabrous and hairy skin were collected. Hairy skin samples were shaved, and underlying tissues (hypodermis, fascia, and muscle) were removed. Tissues were immersion-fixed in 4% paraformaldehyde (PFA) in PBS overnight at 4 °C. Following fixation, tissues were cryoprotected in a graded sucrose series (15%, 20%, and 30% in PBS), embedded in optimal cutting temperature compound (O.C.T.; Thermo Fisher Scientific), snap-frozen on dry ice, and stored at −80 °C. DRG tissues were cryosectioned at 10 μm, spinal cords at 14 μm, and skin at 30 μm using a Leica CM3050S cryostat and mounted on Superfrost Plus slides (Thermo Fisher Scientific) for subsequent staining. These tissue preparations were used for immunohistochemistry (Figures 2A–C, fig. S1A–C) and *in situ* hybridization (Figures 2, fig. S2).

### Immunohistochemistry

Tissue sections were washed in phosphate-buffered saline (PBS) and permeabilized in blocking buffer containing 0.1% Triton X-100, 5% normal donkey or goat serum, and 1% bovine serum albumin (BSA) in PBS for 1 hour at room temperature. Sections were then incubated overnight at 4 °C with the following primary antibodies diluted in blocking buffer:

Rabbit anti-TRPM8 (1:1000; Alomone Labs, ACC-049, lot: ACC049AN1602), Chicken anti-NF200 (1:1000; Abcam, ab72998), Rabbit anti-CGRP (1:1000; Immunostar, 24112) Goat anti-TRPV1 (1:500; Santa Cruz, sc-12498, lot: G2004), Fluorescently conjugated isolectin B4 (IB4; 1:500; Alexa Fluor 488, Thermo Fisher, I21411, lot: 1226926). After three washes in PBS, sections were incubated for 1.5 hours at room temperature with the following species-specific Alexa Fluor–conjugated secondary antibodies (1:1000):Goat anti-Rabbit IgG (H+L), Alexa Fluor 488 (Invitrogen, A11034; lot: 2110499), Goat anti-Rabbit IgG (H+L), Alexa Fluor 555 (Life Technologies, A21429; lot: 2090567), Goat anti-Chicken IgG (H+L), Alexa Fluor 647 (Abcam, ab150171; lot: GR3455191-1), Goat anti-Chicken IgG (H+L), Alexa Fluor 488 (Life Technologies, A11039; lot: 2420700), Donkey anti-Goat IgG (H+L), Alexa Fluor 647 (Abcam, ab150135)

Nuclei were counterstained with DAPI (1:1000 in PBS, 10 minutes; Sigma-Aldrich, 10236276001), followed by final PBS washes. Slides were coverslipped with fluorescence mounting medium (Dako, 11623111).

Imaging was performed using a Zeiss LSM700 laser-scanning confocal microscope or an Olympus CSU-W1 spinning disk confocal system equipped with 20× objectives.

Laser power, exposure time, and acquisition settings were kept constant across all samples. Quantification of marker co-expression, fluorescence intensity, and neuronal counts was conducted using FIJI/ImageJ (NIH) and QuPath (version 0.4.0), with standardized thresholding applied to all images. Neuronal quantification was performed on ≥3 animals per species, with a minimum of 2,000 neurons analyzed per animal. Image analysis was performed blinded to species and condition.

### Immunocytochemistry on cultured cells

Cells were fixed with 4% paraformaldehyde (PFA) in PBS for 10 minutes at room temperature, followed by three PBS washes. Blocking was performed for 1 hour at room temperature using 5% horse serum in PBS, either with or without 0.1% Triton X-100 depending on whether surface or intracellular epitopes were targeted.

Cells were incubated overnight at 4 °C with rabbit anti-TRPM8 (1:1000; Alomone Labs, ACC-049, lot: ACC049AN1602) diluted in either 3% horse serum in PBS or 3% horse serum in 0.1% Triton X-100/PBS. After three PBS washes, Goat anti-Rabbit IgG (H+L), Alexa Fluor 555 (Life Technologies, A21429, lot: 2090567), diluted 1:2000 in PBS were applied for 1.5 hours at room temperature. Nuclei were counterstained with DAPI (1:1000 in PBS, 10 minutes; Sigma-Aldrich, 10236276001), followed by three final PBS washes. Coverslips were mounted in fluorescence mounting medium (Dako, 11623111).

Imaging of HEK293T cells was performed on a Zeiss LSM880 confocal microscope equipped with Airyscan super-resolution and a 63× oil immersion objective. Imaging of N2a cells was performed using an Olympus CSU-W1 spinning disk confocal microscope with a 100× oil objective. All acquisition parameters (exposure, laser intensity, gain, and pixel size) were held constant within each experiment.

Quantification of TRPM8 subcellular localization, including speckling patterns and membrane distribution, was performed using FIJI (ImageJ) and ZEN software (Zeiss). Cells were classified as speckled if discrete punctate structures were observed in the cytoplasm or periphery under high-magnification imaging. At least 30 cells per condition were analyzed from ≥3 independent transfections. Image analysis was performed blinded to construct identity.

Speckling analysis was performed using automated intensity-based segmentation in FIJI (ImageJ). Puncta were detected using the “Analyze Particles” function after local thresholding, with parameters optimized for TRPM8 signal. A cell was classified as “speckled” if it contained ≥2 discrete TRPM8⁺ puncta in the cytoplasm or periphery.

### Multiplex Fluorescent *In Situ* Hybridization (smFISH)

Detection of *Trpm8* and *Ex-Trpm8* mRNA in DRG sections was performed using the RNAscope Multiplex Fluorescent Assay v2 kit (Advanced Cell Diagnostics, ACD Bio, 323100), following the manufacturer’s protocol. Briefly, 10 μm cryosections were first incubated in PBS for 5 minutes, then allowed to dry at 40 °C for 30 minutes. Endogenous peroxidase activity was quenched by incubating the sections with hydrogen peroxide for 15 minutes at room temperature. After an additional 5-minute wash in 100% ethanol, tissue sections were treated with Protease III (ACD Bio, 322337) for 30 minutes at room temperature. Probes were hybridized for 2 hours at 40 °C in a humidified oven. Subsequent signal amplification steps (Amp1, Amp2, Amp3) were performed for 30 minutes each at 40 °C. Detection was carried out using HRP-labeled channel-specific probes (*Trpm8* (ACD Bio, 537651-C1), *Ex-Trpm8* (ACD Bio, 1280181-C2), followed by TSA-mediated fluorophore binding using Opal 570 and Opal 690 reagents (1:750 dilution; Akoya Biosciences, FP1488001KT and FP1497001KT, respectively). HRP-blocking was performed after each fluorophore step.

For combined RNAscope and immunohistochemistry, sections underwent immunostaining immediately following the final HRP-blocking step. Tissue was blocked in 5% normal donkey or goat serum in PBS with 0.1% Triton X-100 for 1 hour at room temperature, followed by overnight incubation at 4 °C with primary antibodies against CGRP (1:1000; ImmunoStar, 24112), TRPV1 (1:500; Santa Cruz Biotechnology, sc-12498, lot: G2004), NF200 (1:1000; Abcam, ab72998), or Alexa Fluor–conjugated IB4 (IB4, 1:500; Thermo Fisher Scientific, I21411, lot: 1226926). After three PBS washes, Alexa Fluor–conjugated secondary antibodies were applied for 1.5 hours at room temperature. Sections were counterstained with DAPI (1:1000 in PBS, 10 minutes; Sigma-Aldrich, 10236276001) and mounted using fluorescence mounting medium (Dako, 11623111).

Imaging was performed using an Olympus CSU-W1 spinning disk confocal microscope with a 20x objective. All imaging parameters (exposure, laser power, contrast) were kept constant across samples. In selected experiments, combined mRNA and protein localization was assessed. Quantification was performed across multiple sections per animal, with >2000 neurons analyzed per group, using tissues from at least five naked mole-rats per group.

### TRPM8 Constructs

A total of five TRPM8 expression plasmids were used in this study. Four constructs were generated in Berlin using a modified pRK9-IRES-GFP vector, and one construct was obtained in Spain containing an IRES-mCherry cassette. All constructs were sequence-verified by Sanger sequencing and used for cell-based calcium imaging and localization assays. Transfected cells were identified based on GFP or mCherry fluorescence.

The Berlin GFP constructs included: MmTRPM8-GFP: full-length mouse TRPM8, HgTRPM8-GFP: full-length naked mole-rat TRPM8 including the extended N-terminal domain, ΔN1–71 HgTRPM8-GFP: truncated HgTRPM8 lacking the first 71 amino acids, HgNter–MmTRPM8-GFP: chimeric construct containing the naked mole-rat N-terminal domain fused to the mouse TRPM8 backbone. The fifth construct, MmTRPM8-mCherry, was generated in Spain and used in competition experiments.

### Cell culture and transfection

HEK293, HEK293T, and Neuro-2a (N2a) cells were maintained at 37 °C in a humidified incubator with 5% CO₂. HEK293 cells (ECACC Cat# 85120602, RRID: CVCL_0045) were cultured either in DMEM-GlutaMAX (Gibco, Thermo Fisher Scientific) supplemented with 10% fetal bovine serum (FBS; PAN Biotech) and 1% penicillin-streptomycin (P/S; Sigma-Aldrich), or in Minimum Essential Medium (MEM; Gibco, 11095-080) supplemented with 10% FBS (Gibco, 16000-044), 1% MEM vitamin solution (Gibco, 11120-037), 100 U/mL penicillin, and 100 μg/mL streptomycin (Gibco, PL15140122), depending on the experimental paradigm. HEK293T cells were maintained under identical conditions as the DMEM group. N2a cells were cultured in DMEM-GlutaMAX supplemented with 45% Opti-MEM (Gibco), 10% FBS, and 1% P/S.

For all experiments, cells were seeded onto 12-mm glass coverslips pre-coated with 0.01% poly-L-lysine (Sigma-Aldrich) and allowed to adhere overnight prior to transfection. HEK293 cells were plated in 6-well plates at a density of 1.5 × 10 cells per well for electrophysiological studies, while similar plating densities were used for imaging. Transfections were carried out using Lipofectamine 2000 (Thermo Fisher Scientific or Invitrogen), following the manufacturer’s instructions. Specifically, 3 μL of Lipofectamine 2000 was mixed with 1 μg of plasmid DNA in 300 μL of Opti-MEM (Thermo Fisher Scientific), incubated for 20 minutes, and applied to the cells for 4 hours. For HEK293T and N2a cells, FuGene 6 (Promega) was used according to the manufacturer’s protocol.

The plasmids used included: MmTRPM8 (mouse TRPM8), HgTRPM8 (full-length naked mole-rat TRPM8 with N-terminal extension), ΔN1–71 HgTRPM8 (truncated HgTRPM8 lacking the first 71 amino acids), HgNter–MmTRPM8 (chimeric construct containing the naked mole-rat N-terminus fused to the mouse TRPM8 backbone), MmTRPM8-mCherry (mouse TRPM8). All constructs encoded either canonical or isoform-specific variants of TRPM8, with or without fluorescent reporters (GFP or mCherry). A total of 0.25 μg DNA per well was used for all conditions.

Electrophysiological recordings were conducted 24–36 hours post-transfection. For calcium imaging and immunocytochemistry, cells were imaged 24–48 hours post-transfection. For each condition, data were collected from at least three independent transfections. Quantification was performed across ≥30 cells per condition and pooled from at least three biological replicates.

### Calcium imaging in HEK293 cells

Ratiometric calcium imaging was performed using the fluorescent calcium indicator Fura-2 (Invitrogen). HEK293 cells were incubated with 5 μM Fura-2 AM and 0.02% Pluronic F-127 (prepared from a 200 mg/mL DMSO stock) in HEPES-buffered saline for 45 minutes at 37 °C, followed by a 15-minute de-esterification at room temperature. Coverslips were mounted in a perfusion chamber on a Leica DMIRES-2 inverted microscope equipped with a Lambda 421 LED light source (Sutter Instruments) and a Retiga R6 CCD camera (QImaging).

Excitation wavelengths alternated between 340 and 380 nm, and emission was collected at 510 nm. Images were acquired every 2–3 seconds using MetaFluor software (Molecular Devices). GFP⁺ or mCherry⁺ transfected cells were identified and manually segmented using ROI tools. Calcium responses were expressed as changes in the 340/380 fluorescence ratio (ΔFura-2).

Extracellular solution (Buffer) was (in mM): 140 NaCl, 3 KCl,1.3 MgCl_2_, 10 HEPES, and 10 glucose, 2.4 CaCl_2_ (290 mOsm/L, pH adjusted to 7.4 with NaOH. Recordings were performed at a basal temperature of 33 ± 1 °C. Cells were sequentially stimulated with the following conditions: warm buffer (∼40 °C), cold buffer (16 °C), menthol (200 μM), icilin (10 μM), and combined icilin + cold. Each stimulus was applied for 60– 90 seconds, with 2-minute washout intervals between conditions. All recordings were conducted under continuous perfusion. A response was considered positive if the ΔFura-2 ratio increased by >0.08 above baseline, based on thresholds determined from vehicle control recordings.

### Electrophysiology in cultured cells

Whole-cell voltage-clamp recordings were performed on transiently transfected HEK293 cells using borosilicate glass patch pipettes (Sutter Instruments) with a resistance of 4–8 MΩ. Recordings were conducted concurrently with temperature measurements. Electrical signals were amplified with an Axopatch 200B patch-clamp amplifier (Molecular Devices) and digitized using a Digidata 1440A interface (Molecular Devices). Stimulus application and data acquisition were controlled with pCLAMP 11 software (Molecular Devices).

For electrophysiological experiments in HEK293 cells, extracellular solution was (in mM): 140 NaCl, 3 KCl,1.3 MgCl_2_, 10 HEPES, and 10 glucose, 2.4 CaCl_2_ (290 mOsm/L, pH adjusted to 7.4 with NaOH). The intracellular solution was (in mM): 135 CsCl, 2 MgCl2, 10 HEPES, 1 EGTA, 5 Na2ATP and 0.1 NaGTP, adjusted to pH 7.4 with CsOH (280 mOsm/L). Recordings were performed at a basal temperature of 33 ± 1C. The chemical agonists used are: icilin (Sigma, I9532, stock 10 mM in DMSO), and menthol (Scharlau, stock 300 mM in ethanol)

Glass coverslips (6 mm diameter) containing cultured cells were placed in a microchamber and continuously perfused with solution maintained at 33 ± 1 °C. Temperature control was achieved using a Peltier device (CoolSolutions, Cork, Ireland) positioned at the chamber inlet and regulated via a feedback system as described by Reid^66^. Cold sensitivity was assessed by reducing the bath temperature to approximately 18 °C. Temperature was monitored using an IT-18 T-type thermocouple connected to a BAT-12 microprobe thermometer (Physitemp Instruments), with the signal digitized through an Axon Digidata 1440A A/D converter and recorded using Clampex 11 software (Molecular Devices).

Once a Gigaohm seal was achieved and the whole-cell configuration established, cells were voltage-clamped at a holding potential of –60 mV. To assess the effects of TRPM8 agonists (cold, icilin 10 μM, and menthol 200 μM) on the various plasmid constructs, current responses were elicited by repetitive voltage ramps (–110 to +180 mV, 400 ms duration) delivered at 0.33 Hz. Current amplitudes were measured at –100 mV and +100 mV, and peak amplitudes were compared to those recorded under control conditions.

Data are normalized to membrane capacitance (pA/pF) and reported as mean ± SEM. For multiple comparisons of means, one-way ANOVAs were performed, followed by Bonferroni’s post-hoc analysis, using GraphPadPrism version 8.00 for Windows (GraphPad Software). During statistical analysis, normality of the data distribution was determined with the Shapiro–Wilk test.

To evaluate shifts in the voltage dependence of TRPM8 activation during exposure to cold or chemical agonists, the I–V curves to the –110 to +180 mV voltage ramps were fitted using the BoltzIV function in OriginPro 2024 (OriginLab Corporation, Northampton, MA, USA) as described previously^67^. Parameters extracted included Gmin and Gmax (minimum and maximum whole-cell conductance), Erev (current reversal potential), V_1/2_ (voltage at half-maximal activation), and dx (slope factor). We determined the Erev parameter independently for each condition by measuring the average voltage at which the current reversed sign, which was then kept constant during the fitting process. All fittings were carried out with the Levenberg–Marquardt method implemented in the OriginPro version 2024 software.

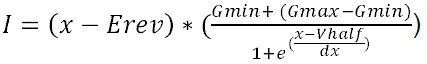

### DRG neuron culture

Lumbar dorsal root ganglia (DRGs; L3–L5) were isolated from adult mice and naked mole-rats and transferred into cold plating medium consisting of DMEM/F-12 (Gibco, Thermo Fisher Scientific) supplemented with 10% fetal horse serum (FHS; Life Technologies) and 1% penicillin–streptomycin (P/S; Sigma-Aldrich). Tissues were enzymatically digested in 1.25% Collagenase IV (1 mg/mL; Sigma-Aldrich) for 45 minutes at 37 °C, followed by 2.5% Trypsin (Sigma-Aldrich) for 15 minutes at 37 °C. After digestion, ganglia were gently triturated using fire-polished Pasteur pipettes to generate a single-cell suspension, which was passed through a 20 μm nylon mesh filter (Millex, Merck Millipore, SLGV004SL). Neurons were further purified using a 15% fraction V BSA gradient (Sigma-Aldrich, A4161-1G) and collected by gentle centrifugation. Neurons were plated onto 8 mm glass coverslips pre-coated with poly-L-lysine (100 μg/mL; Sigma-Aldrich) and laminin (20 μg/mL; Invitrogen) for calcium imaging or immunostaining.

Mouse DRG neurons were cultured overnight at 37 °C in a humidified incubator with 5% CO₂. Naked mole-rat DRG neurons were cultured under identical conditions, except at 32 °C with 5% CO₂.

### Calcium imaging in DRG neurons

Calcium imaging was performed 18–24 hours after plating. DRG neurons were loaded with 5 μM Cal-520 AM (AAT Bioquest) diluted in HEPES-buffered saline, incubated for 45 minutes at 32 °C, and de-esterified for 15 minutes at room temperature. Coverslips were mounted in a temperature-controlled perfusion chamber maintained at 32 °C and imaged on an Olympus BX51WI upright fluorescence microscope equipped with a pE-340fura LED light source (CoolLED) and a CoolSNAP HQ2 CCD camera (Photometrics).

All imaging was conducted under continuous perfusion using a closed-loop inline temperature control system (Cool Solutions Heat/Cooled Temperature Control System, Autom8, UK), which actively regulated perfusate temperature via integrated heating and cooling modules. The perfusion line was connected to a temperature feedback controller that maintained a stable chamber temperature of 32 °C throughout the experiment.

Neurons were sequentially stimulated with a series of thermal and chemical stimuli, with the exact order and combination adjusted according to the experimental condition. Standard protocols typically included: warm buffer (∼32 °C), cold buffer (16 °C), menthol (200 μM; Sigma, Cat# M2772), AMTB (10 μM; Biotechne, Cat# 3989/50) + cold, cold alone, and potassium chloride (50 mM). For some experiments, cold was applied either before or after pharmacological agents depending on the specific treatment paradigm. For pharmacological inhibition assays, neurons were pre-incubated with AMTB (10 μM) or PBMC (10 μM; Focus Biomolecules, Cat# 10-1413) for 60 minutes prior to cold stimulation. In these cases, cold responsiveness was assessed either in the continuous presence of the antagonist or following pre-treatment, depending on the protocol. All stimuli were washed out thoroughly between applications to minimize carry-over. Neurons that failed to respond to the final KCl (50 mM) application were excluded from analysis.

Fluorescence signals were acquired at 0.5 Hz using MetaFluor software (Molecular Devices), and calcium responses were quantified as changes in fluorescence intensity over baseline (ΔF/F₀). Neurons were classified as cold-sensitive if they exhibited a response amplitude >10% (ΔF/F₀ > 0.10) following cold stimulation. Data were collected from at least three animals per group, with ≥200 neurons analyzed per condition.

### *Ex vivo* skin–nerve electrophysiology

Electrophysiological recordings were performed on tissues from 21 mice and 17 naked mole-rats. Hairy skin from the hind paw, along with the saphenous nerve, was dissected and used in an *ex vivo* single-fiber skin–nerve preparation, as previously described^1,2,41^. Following euthanasia (cervical dislocation for mice, decapitation for naked mole-rats), skin–nerve preparations were transferred to a two-chamber organ bath. The skin was pinned epidermal side up (“outside-out” configuration) and superfused with oxygenated (95% O₂, 5% CO₂) synthetic interstitial fluid (SIF) maintained at 32 °C. The SIF composition was (in mM): 123 NaCl, 3.5 KCl, 0.7 MgSO₄, 1.7 NaH₂PO₄, 2.0 CaCl₂, 9.5 sodium gluconate, 5.5 glucose, 7.5 sucrose, and 10 HEPES; pH adjusted to 7.4.

The saphenous nerve was placed in a neighboring mineral oil–filled chamber, and the perineurium was carefully removed under a dissection microscope. Fine nerve filaments were teased and placed on a platinum wire electrode for extracellular recording.

Sensory fibers were identified using mechanical stimulation (blunt probes, glass rods) and electrical search stimuli delivered to the receptive field via a concentric bipolar electrode connected to an isolated stimulator (Digitimer Ltd., UK). Conduction velocity (CV) was calculated from spike latency and physical distance using the formula: CV = distance / latency. Based on CV, fibers were classified as Aβ (>10 m/s), Aδ (1.5– 10 m/s), or C (<1.5 m/s).

Cold sensitivity was assessed by superfusing the receptive field with SIF cooled to ∼10 °C via gravity flow. Spike activity was recorded before and after bath application of the TRPM8 antagonist AMTB (10 μM), perfused for at least 10 minutes. Fibers were categorized as TRPM8-dependent if cold-evoked spike frequency was abolished or reduced by ≥80% following AMTB application.

Recordings were acquired using a PowerLab 4/30 system (ADInstruments) and analyzed with LabChart 7.1 and the Spike Histogram extension. Only fibers with stable receptive fields, clearly identifiable action potentials, and reliable conduction velocity were included. Spike quantification and TRPM8-dependence classification were performed using semi-automated detection, followed by manual confirmation. Analyses were conducted blinded to species identity. Instantaneous firing rate (IFR) was calculated as the inverse of the inter-spike interval (ISI), using the formula: IFR(t) = 1 / ISI(t).

### RNA-Seq Analysis and Cross-Species Sequence Alignment

Naked mole-rat dorsal root ganglia (DRG) RNA sequence data was obtained from Eigenbrod et al., 2019^30^. Reads were cleaned using Trimmomatic^68^, part of the transcriptome assembly package Trinity^69^. Trimmed reads were mapped to the recently annotated^70^. Ensembl naked mole-rat genome (GCA_944319715.1), using STAR (v2.7.10a,^71^). Aligned reads were sorted and indexed using SAMtools (v1.17,^72^). Exon read coverage was calculated with featureCounts (v2.0.6)^73^ using per exon counts (–f) and allowing for multi-mapping reads (–M) and overlapping features (–O). Plotting of the data was done using Python and matplotlib (v3.7).

Sequences and exon structure of the orthologs from the human, guinea pig, mouse and clawed frog were obtained from the National Center for Biotechnology Information (NCBI) database. Sequence alignments were performed using MAFFT^74^. Visualization of the naked mole-rat exon structure and read mapping was done with integrative genomics viewer^75^.

### Statistical Analysis

Quantification of cell diameter, marker co-localization, and subcellular speckling intensity was performed using FIJI/ImageJ (NIH) and QuPath (v0.4.3). All image-based analyses were conducted using standardized thresholding parameters, and where applicable, analyses were performed blinded to experimental condition.

Statistical analyses were conducted using GraphPad Prism (v9), R (v4.4.3), and Origin (OriginLab, 2024b), depending on dataset type and experimental design. Data normality was assessed using the Shapiro–Wilk test. For parametric data, one-way ANOVA was applied, followed by Tukey’s multiple comparisons tests for all pairwise group comparisons. Unpaired two-tailed Student’s t-tests were used for pairwise comparisons. Specific statistical tests used, including exact n, p-values, and correction methods, are provided in the corresponding figure legends. All data are presented as mean ± SEM unless otherwise specified. Statistical significance was defined as *p* < 0.05. The following significance notation was used: *p* < 0.05 (*), *p* < 0.01 (**), *p* < 0.001 (***), and *p* ≥ 0.05 (ns).

## Acknowledgements

This work was supported by the European Research Council (ERC-2023-ADG-101142335, JFAP) and (ERC-2018-ADG-789128 to GRL), the Deutsche Forschungsgemeinschaft (SFB 1315, SFB TRR 384, JFAP), and the Human Frontier Science Program RGP023/2024, Generalitat Valenciana CIGRIS/2022/056 to FV, Spanish State Research Agency project PID2022140961OB-100 to FV and the “Severo Ochoa” Center of Excellence grant to the IN (CEX2021-001165-S to FV),and was co-financed by the European Regional Development Fund (ERDF). LW was supported by the fellowship from the Chinese Scholarship Council. We thank members of the Lewin lab for their comments on the MS. We thank Maria Braunschweig for excellent technical assistance and Anika Mühlenberg and Gabriela Pflanz for naked mole-rat husbandry.

## Author Contributions

Conceptualisation, L.W., G.R.L, F.V., V.B, and J.F.A.P; methodology, G.R.L, L.W, F.V, EP; formal analyses, L.W., F.V., A.R., E.P., V.B; investigation, LW; A.R, AN, F.V., W.L., D.M.A., J.J.; writing original draft preparation, L.W, V.B. and G.R.L.; resources G.R.L. V.B., J.F.A.P.; funding, G.R.L, F.V. and J.F.A.P; supervision, V.B., G.R.L, F.V., E.P. and J.F.A.P.; Visualization L.W., F.V., D.M.A, V.B., G.R.L.

## Supplementary Figures

**Figure S1.**
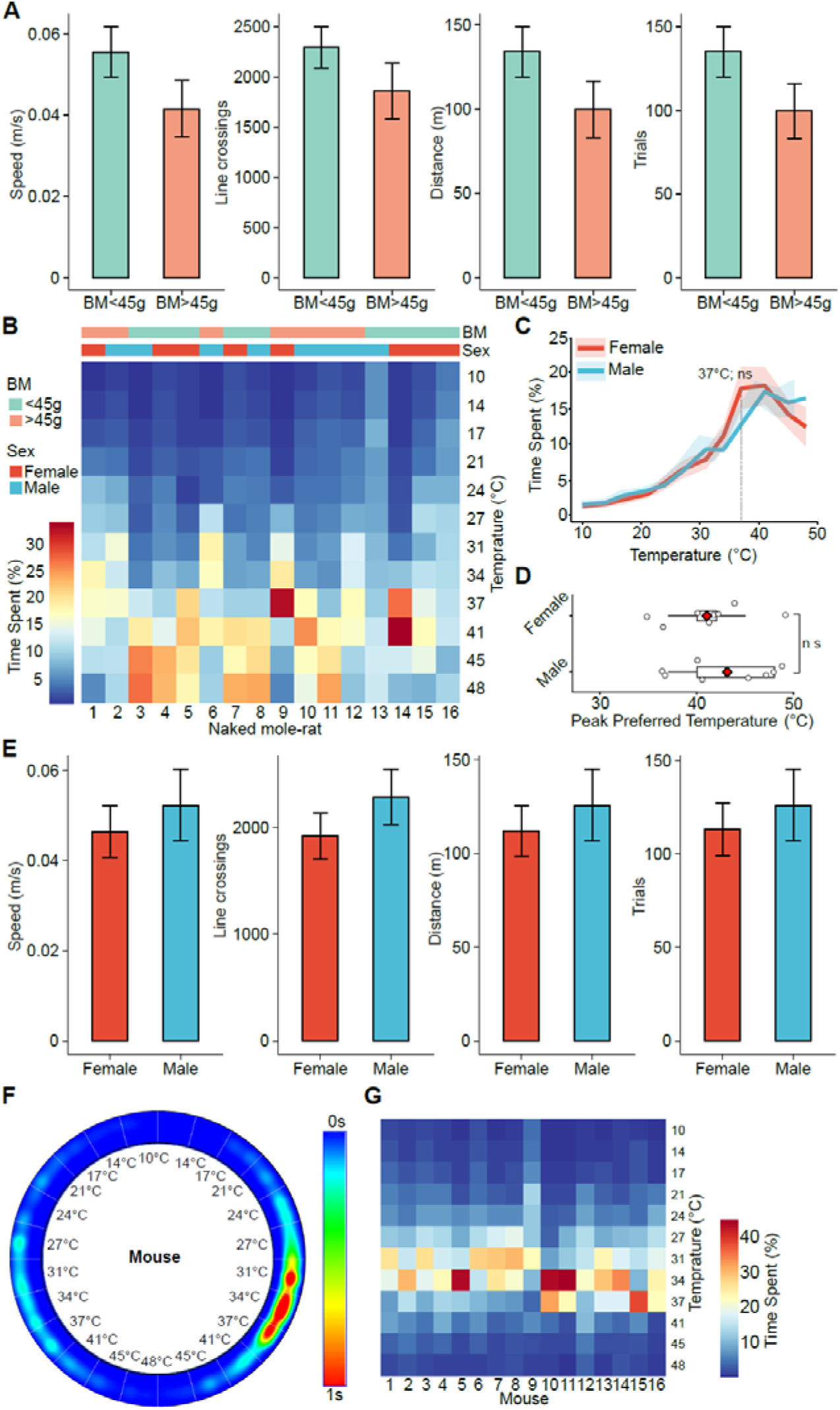
Warm seeking is not different between naked mole-rat genders. (**A**) Summary locomotor metrics including speed, line crossings, total distance, and trial count, stratified by body mass (<45 g vs. >45 g). Data are presented as mean ± SEM. Unpaired two-tailed Student’s t-test, ns. (**B**) Heatmap of temperature zone occupancy for individual animals during the thermal gradient assay, annotated by body mass (BM) and sex. (**C**) Time spent per temperature bin, grouped by sex. *n* = 8 females, *n* = 8 males. Shaded areas indicate mean ± SEM. Unpaired two-tailed Student’s t-test at peak zone, *p* = 0.15, ns. (**D**) Peak preferred temperature per animal, grouped by sex. Red diamonds indicate group means. Unpaired two-tailed Student’s t-test, ns. **(E**) Locomotor parameters by sex (speed, line crossings, distance, and trial entries). Data shown as mean ± SEM. Unpaired two-tailed Student’s t-tests, all comparisons ns. (**F**) Representative occupancy heatmap of pooled data from all mice (n = 16; 8 male, 8 female) over 40 minutes in a 12-zone thermal gradient ring. (**G**) Heatmap of temperature zone occupancy for individual mouse during the thermal gradient assay.

**Figure S2.**
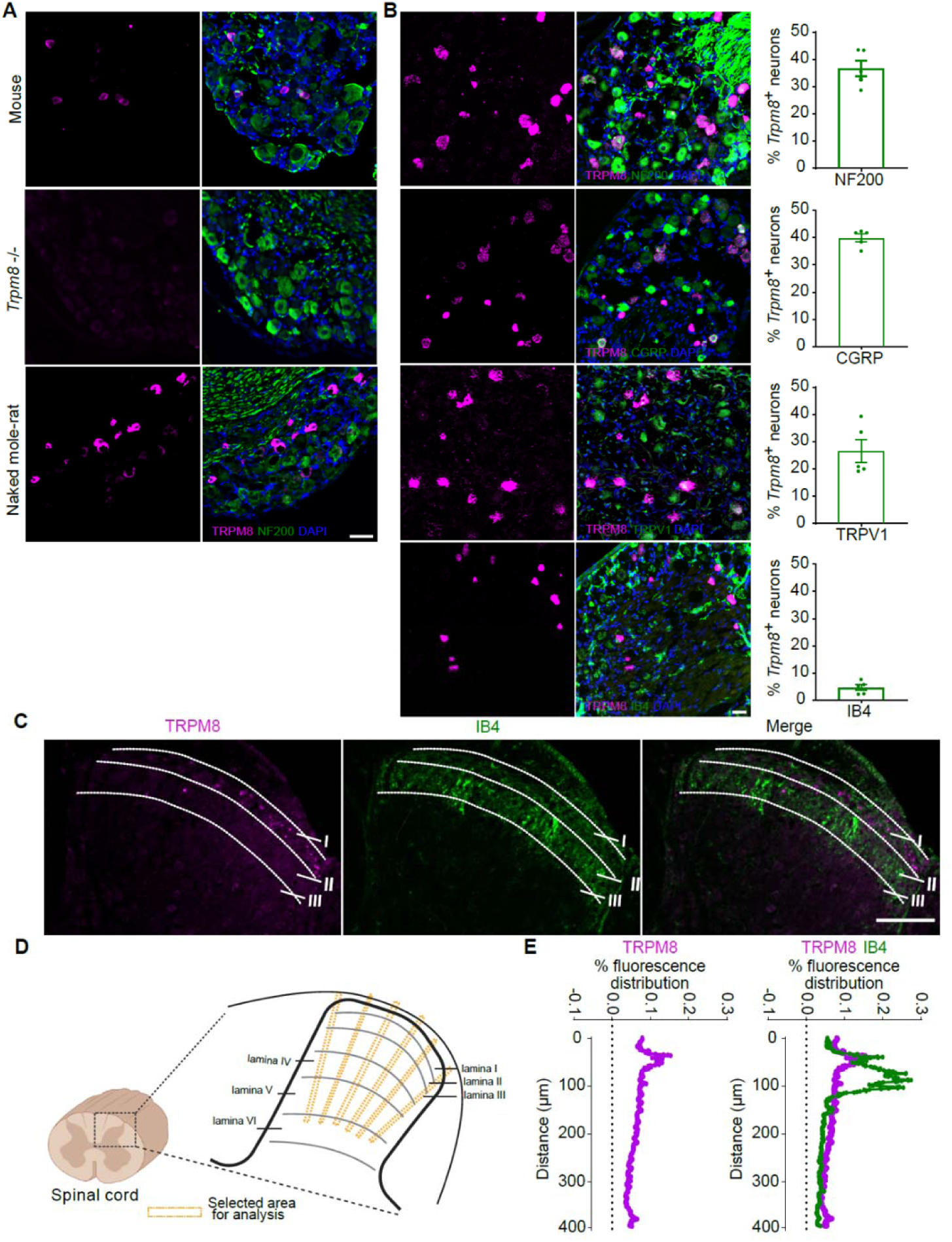
Comparative analysis of TRPM8 expression in naked mole-rats and mice. **(A)** Immunofluorescence analysis of TRPM8 expression in dorsal root ganglia (DRGs) of C57BL/6 mice (top), Trpm8^⁻/⁻^ mice (middle), and naked mole-rats (bottom). Sections were stained for TRPM8 (magenta), NF200 (green; myelinated neurons), and DAPI (blue). Scale bar: 50 μm. **(B)** Cellular identity of *Trpm8* mRNA-expressing neurons in naked mole-rat DRGs assessed by RNAScope fluorescent in situ hybridization. Representative images show *Trpm8* mRNA (magenta) co-expression with NF200, CGRP (peptidergic marker), TRPV1, or IB4 (non-peptidergic marker) (green). Quantification (right) shows the percentage of *Trpm8*⁺ neurons expressing each marker. Data represent mean ± SEM from *n* = 5 animals, >2000 neurons analyzed per condition. Scale bar: 50 μm. **(C)** Distribution of TRPM8⁺ primary afferent terminals in the lumbar spinal cord of naked mole-rats. Immunostaining for TRPM8 (magenta) and IB4 (green) reveals innervation patterns in the dorsal horn. White dashed lines delineate laminar boundaries. Scale bar: 100 μm. **(D)** Schematic illustration of spinal cord laminar organization and analysis strategy. The boxed region (left) indicates the dorsal horn area selected for quantification of TRPM8⁺ primary afferent terminals, focusing on laminae I–III (right), which receive direct input from cold-sensitive sensory neurons. **(E)** Spatial distribution of TRPM8 fluorescence within the dorsal horn. Left: fluorescence intensity profile of TRPM8⁺ primary afferent terminals across laminae I–VI, normalized to the total TRPM8 signal within the selected region of interest (as defined in D). Right: overlay of TRPM8 and IB4 fluorescence distributions, each normalized to its respective total signal within the same region to correct for inter-sample variation in staining and image acquisition. TRPM8⁺ terminals are primarily concentrated in lamina I, consistent with their role in transmitting cold sensory input to superficial spinal layers.

**Figure S3.**
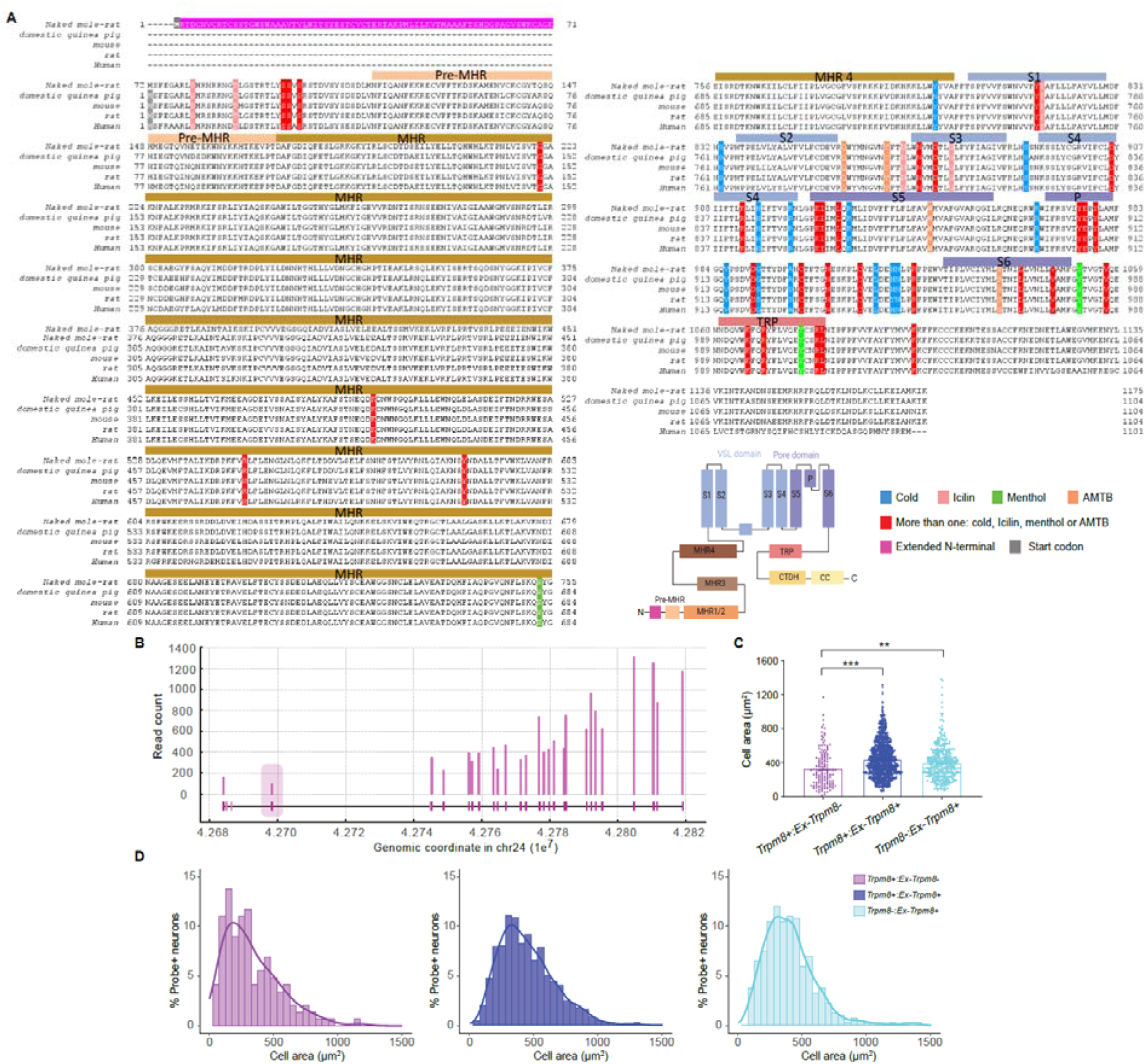
Evolutionary conservation and expression of extended TRPM8 isoform in naked mole-rats. **(A)** Multiple sequence alignment of TRPM8 orthologs. Protein sequences from naked mole-rats, guinea pigs, and other mammals were aligned to identify conserved domains and residues. The extended N-terminal domain unique to naked mole-rats is highlighted in magenta. Conserved structural domains, including the pre-MHR and melastatin homology regions (MHR1–4), the transmembrane domain, and the pore-forming region, are annotated. Functionally important residues mediating responses to cold, icilin, menthol, and interactions with AMTB are color-coded. Right: schematic showing domain architecture of extended TRPM8 isoforms. **(B)** Naked mole-rat specific exon shows moderate expression, confirming naked mole-rat-specific exon transcriptional activity. Read coverage for each exon is shown and it’s genomic position. DRG RNAseq data for all isoforms was included. Exon structure is shown below, with non-expressed predicted exons in pale colour. The magenta colored box marks the first coding exon, containing the naked mole-rat specific sequence. **(C)** Quantification of soma size in neurons expressing different TRPM8 isoforms. Violin plots show the distribution of cell body areas among neurons expressing *Trpm8* only, *Ex-Trpm8* only, or both isoforms. Horizontal bars indicate medians. Statistical comparisons were performed using one-way ANOVA with post hoc tests; \*\**p* < 0.01, \*\*\**p* < 0.001. (D) Histogram plots showing cell area distributions for neurons expressing each TRPM8 isoform type. Each panel corresponds to a different group: *Trpm8*⁺:*Ex-Trpm8*⁻ (left), *Trpm8*⁻:*Ex-Trpm8*⁺ (middle), and *Trpm8*⁺:*Ex-Trpm8*⁺ (right). Gaussian curve fits are overlaid for visualization.

**Figure S4.**
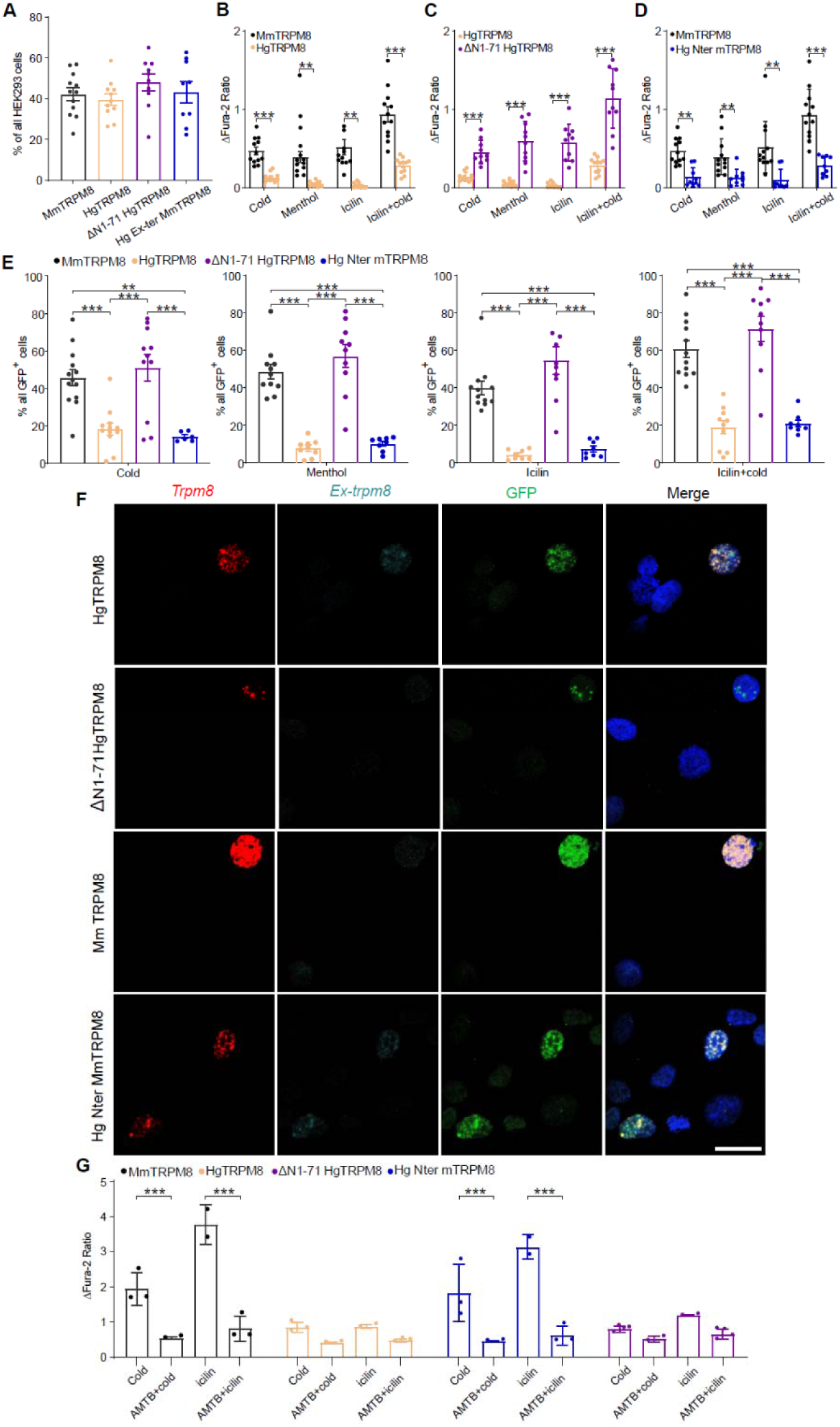
TRPM8 isoform-dependent calcium signals in HEK293 cells. **(A)** Percentage of GFP⁺ HEK293 cells following transfection with the indicated TRPM8 constructs, representing transfection efficiency. **(B–D)** Quantification of ΔFura-2 ratio in response to cold (18°C), menthol (200 μM), icilin (10 μM), and combined icilin + cold stimulation. **(B)** MmTRPM8 vs. full-length naked mole-rat TRPM8 (HgTRPM8). **(C)** HgTRPM8 vs. N-terminal deletion construct (ΔN1–71 HgTRPM8). **(D)** MmTRPM8 vs. chimeric construct containing the naked mole-rat N-terminal extension (HgNter–MmTRPM8). Data shown as mean ± SEM. One-way ANOVA with Tukey’s post hoc test; \*\**p* < 0.01, \*\*\**p* < 0.001. **(E)** Percentage of GFP⁺ cells responding to each stimulus across TRPM8 constructs. One-way ANOVA with Tukey’s post hoc test; \*\*\**p* < 0.001. **(F)** Representative RNAScope images showing *Trpm8* (red) and *Ex-Trpm8* (cyan) transcripts in GFP-labeled (green) HEK293T cells expressing each TRPM8 construct. Nuclei stained with DAPI (blue). Scale bar: 30 μm. **(G)** ΔFura-2 responses to cold and icilin in the presence or absence of the TRPM8 antagonist AMTB (10 μM), across all four constructs. Data represent mean ± SEM. One-way ANOVA with Tukey’s post hoc test; \*\*\**p* < 0.001.

**Figure S5.**
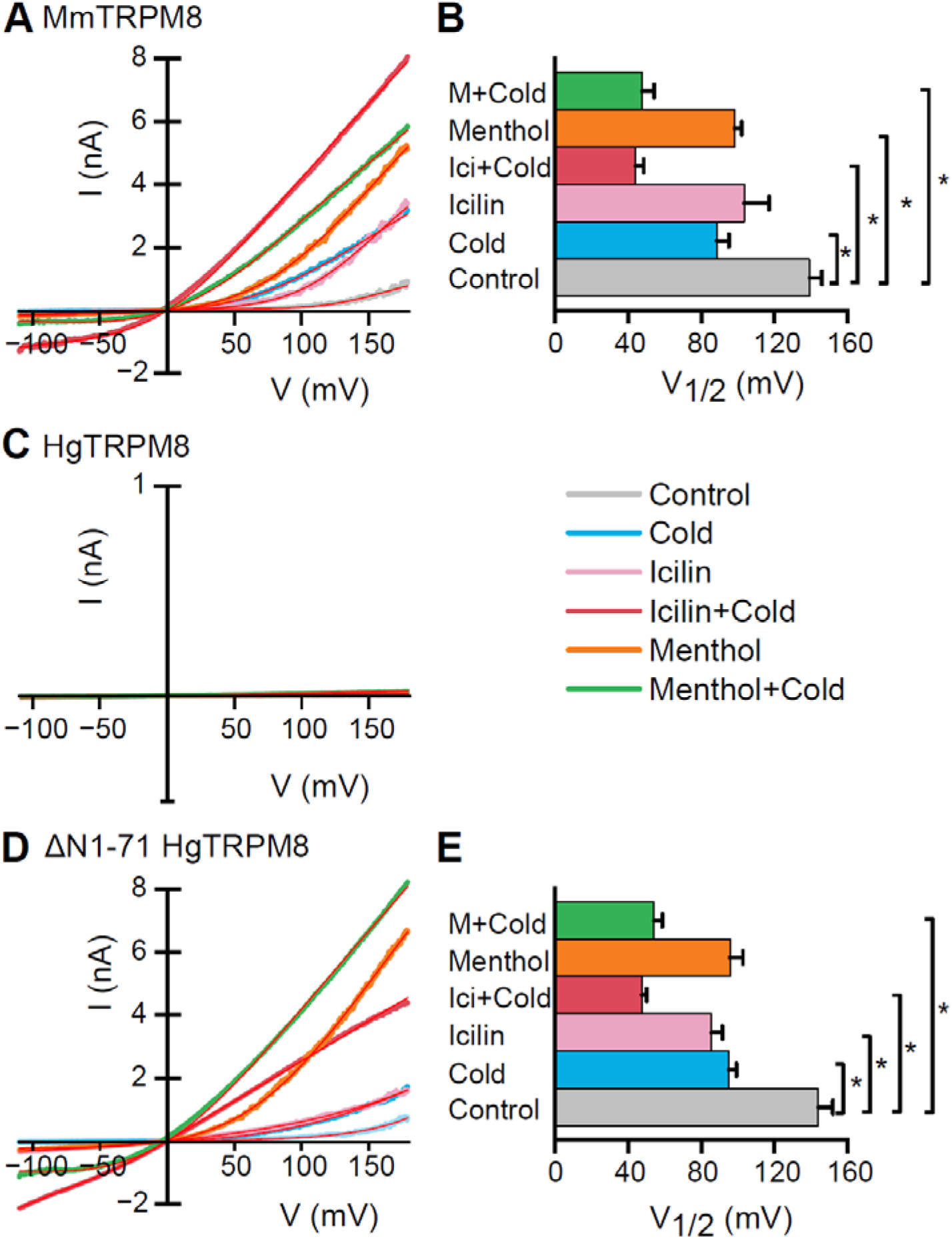
N-terminal deletion restores the activation properties of naked mole-rat TRPM8 channels. **(A)** Whole-cell ramp I–V (–110 to 180 mV) relationships recorded sequentially from the same HEK293 cell expressing mTRPM8 under different experimental conditions: control (33 °C), cold (∼ 16 °C), icilin (10 µM), icilin + cold, menthol (100 µM), menthol + cold. The red lines, superimposed on each trace, represent the fit of the current to a linearized Boltzmann function (see Materials and Methods). **(B)** Summary histogram (mean ± SEM, n = 6) of the midpoint of voltage activation (V_1/2_) of TRPM8 channels under different experimental conditions. **(C)** Expression on naked mole-rat TRPM8 (HgTRPM8) produced minimal currents that could not be properly fitted. Representative example of n = 7 recordings. Note the change in current scale. **(D)** Deletion of the N-terminus of naked mole-rat TRPM8 (ΔN1-71 Hg TRPM8) restored the activation properties of the TRPM8 current, yielding similar V_1/2_ values (n = 6). **(E)** Statistical significance was assessed by one-way ANOVA followed by Tukey’s post hoc test.

**Figure S6.**
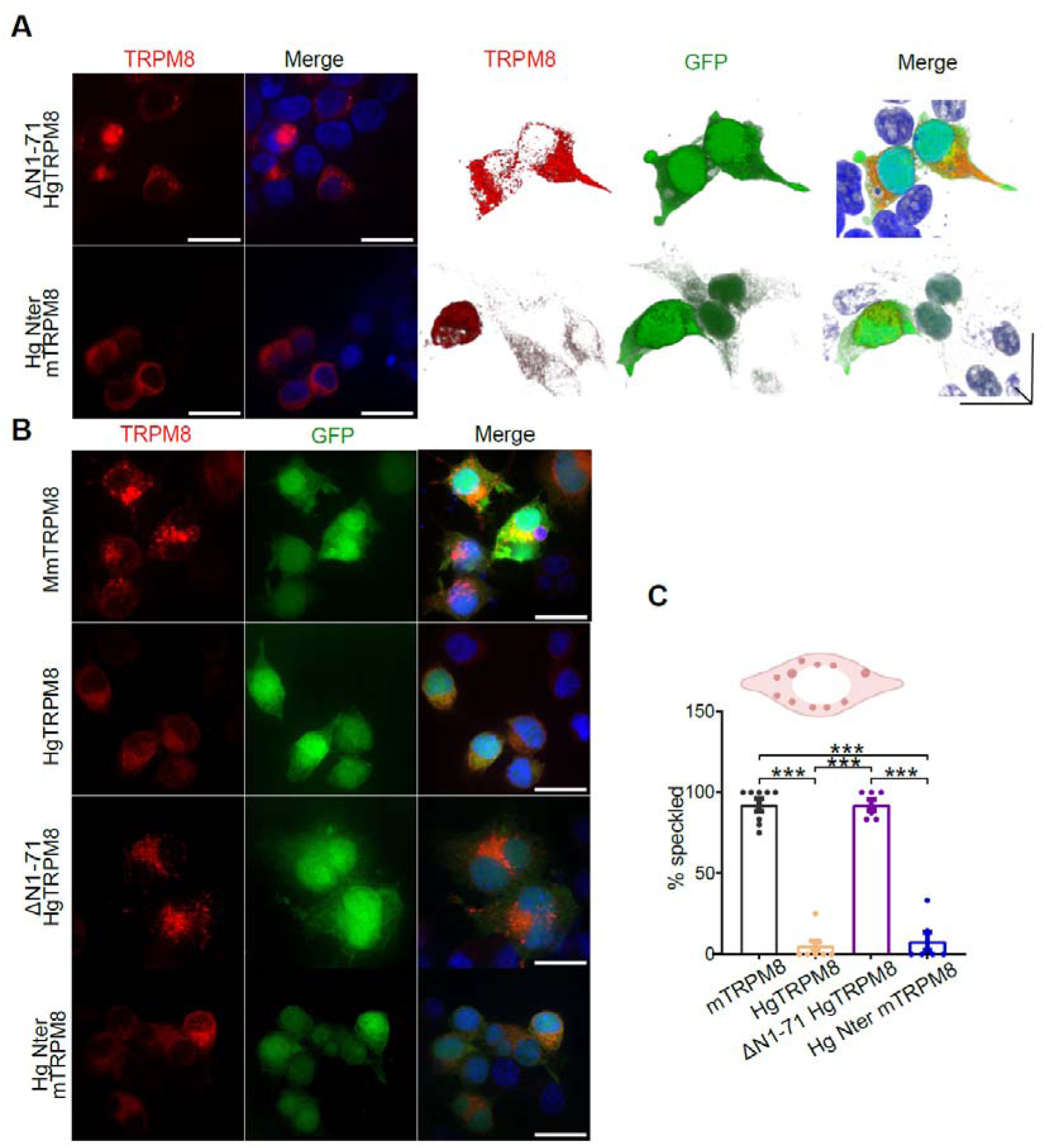
Subcellular localization of TRPM8 isoforms in HEK293T and N2a cells. **(A)** Immunofluorescence images of HEK293T cells transfected with ΔN1–71 HgTRPM8 or HgNter– MmTRPM8. Cells were stained for TRPM8 (red), GFP (green), and nuclei (DAPI, blue). Scale bars: 20 μm. **(B)** Immunofluorescence staining of N2a cells transfected with different TRPM8 isoforms. Images show TRPM8 (red), GFP (green), and nuclei (DAPI, blue). Scale bars: 20 μm. **(C)** Quantification of TRPM8⁺ cells displaying speckled staining patterns in N2a cells. Speckled cells were defined as those containing ≥2 discrete TRPM8⁺ puncta, as determined by automated segmentation in FIJI (see Methods). Data represent mean ± SEM, *n* = 30 cells per construct from ≥3 independentransfections. One-way ANOVA with Tukey’s post hoc test; \**p* < 0.001 for all indicated pairwise comparisons.

**Figure S7.**
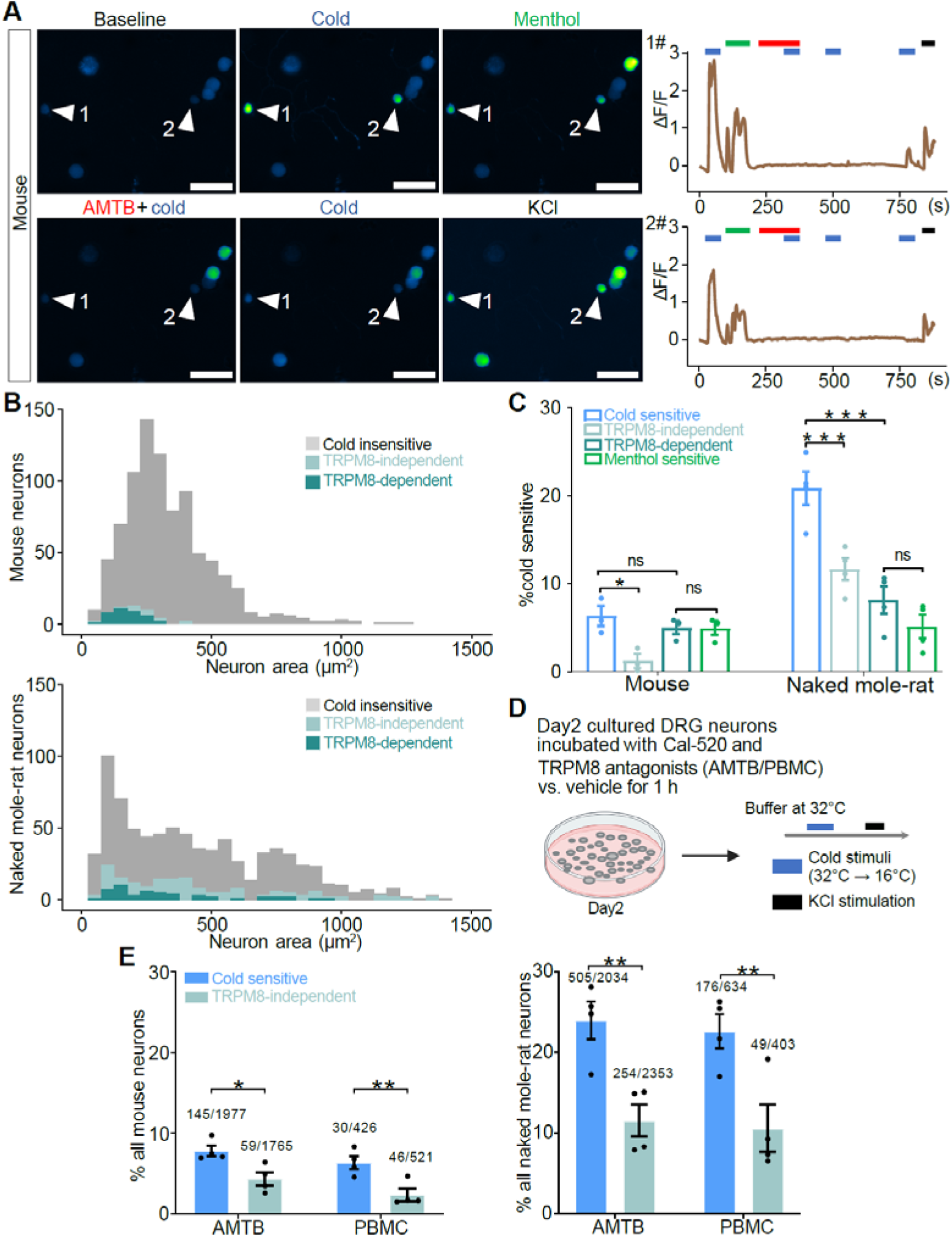
Classification of TRPM8-dependent and TRPM8-independent cold responses in DRG neurons. **(A)** Representative calcium imaging data showing responses of mouse (top) and naked mole-rat (bottom) DRG neurons to cold (32°C → 16°C), menthol (200 μM), AMTB (10 μM) + cold, and KCl (50 mM). Left panels show fluorescence images (Cal-520 signal), and right panels display corresponding ΔF/F₀ traces for two example neurons (1, 2) before and after TRPM8 inhibition. Scale bars: 50 μm. **(B)** Distribution of soma area for cold-insensitive neurons, TRPM8-independent cold-sensitive neurons, and TRPM8-dependent cold-sensitive neurons in mouse (top) and naked mole-rat (bottom) DRG cultures. **(C)** Proportion of cold-sensitive (TRPM8-dependent + TRPM8-independent) and cold-insensitive neurons among all recorded neurons in mouse (*n* = 860 neurons from 3 mice) and naked mole-rat (*n* = 892 neurons from 4 animals) DRGs, using the first calcium imaging protocol. **(D)** Percentage of cold-sensitive neurons in mouse and naked mole-rat DRGs subdivided by TRPM8-dependence and menthol sensitivity. One-way ANOVA with Tukey’s post hoc test; \**p* < 0.05, \*\**p* < 0.01, \*\*\**p* < 0.001, ns = not significant. **(E)** Schematic of the second calcium imaging protocol: DRG neurons were cultured from mouse and naked mole-rat (at 37°C and 32°C, respectively), incubated with Cal-520 and TRPM8 antagonists (AMTB or PBMC) or vehicle for 1 hour on Day 2, and stimulated with cold (32°C → 16°C) and KCl. **(F)** Quantification of cold-sensitive neurons following pharmacological inhibition of TRPM8 using a modified calcium imaging protocol. Percentage of cold-sensitive neurons in each experimental group. For mouse, cold-sensitive neurons were observed in the AMTB vehicle group (145/1977), AMTB-treated group (59/1765), PBMC vehicle group (30/426), and PBMC-treated group (46/521). For naked mole-rat, cold-sensitive neurons were observed in the AMTB vehicle group (505/2034), AMTB-treated group (254/2353), PBMC vehicle group (176/634), and PBMC-treated group (49/403). All treatment and vehicle conditions are shown in shades of cyan, with TRPM8-independent responses indicated in light teal. Data represent recordings from *n* = 4 animals per species. One-way ANOVA; \**p* < 0.05, \*\**p* < 0.01.

**Figure S8.**
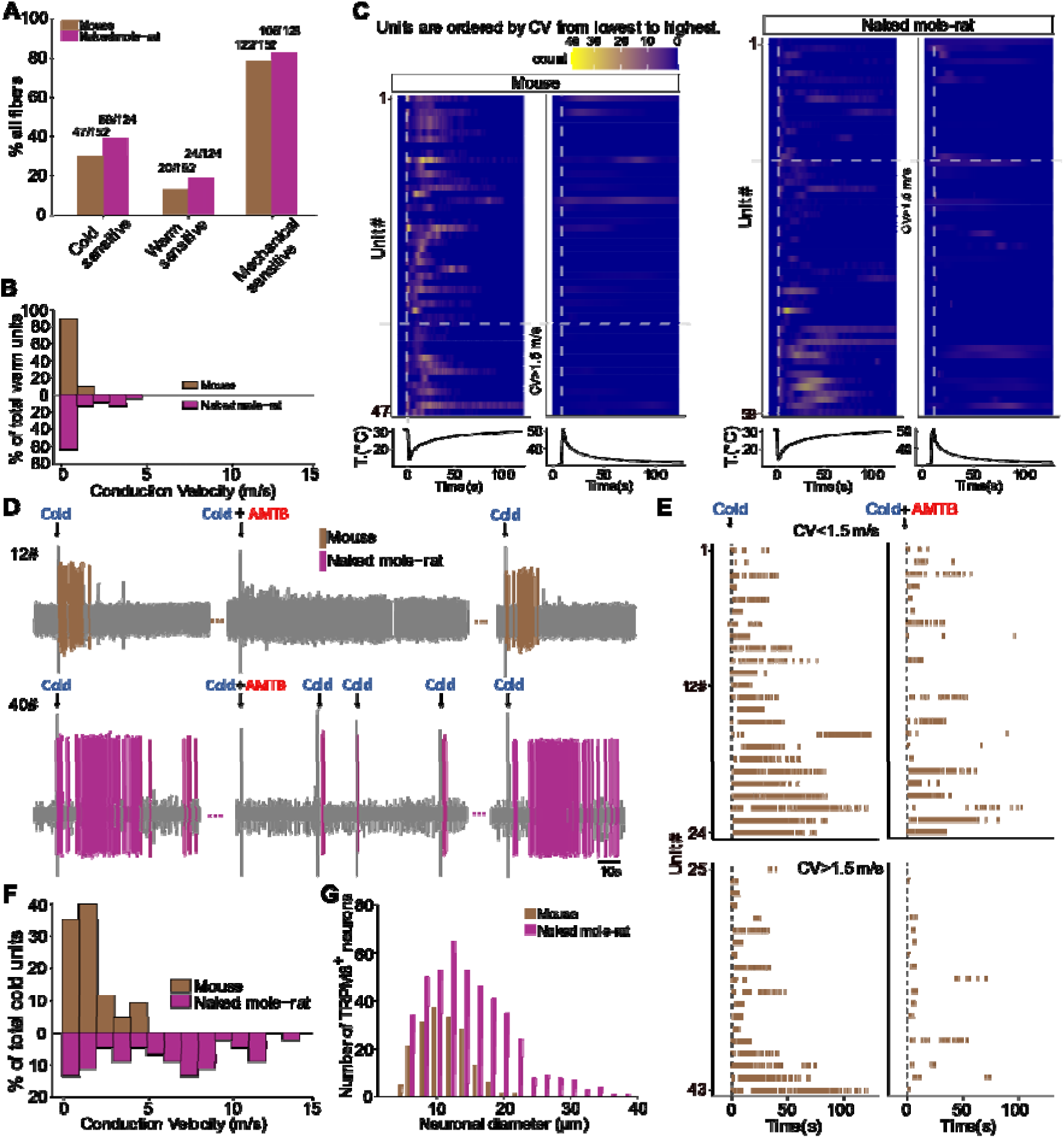
Functional classification and TRPM8-dependence of thermosensitive sensory fibers. **(A)** Proportions of cold-sensitive, warm-sensitive, and mechanically sensitive fibers among all recorded units in mice (152 units from 13 animals) and naked mole-rats (124 units from 11 animals). Counts are indicated above each bar. **(B)** Distribution of conduction velocities among warm-sensitive fibers from mice (*n* = 20 units from 13 animals) and naked mole-rats (*n* = 24 units from 11 animals), units are grouped by conduction velocity (CV > 1.5Lm/s). **(C)** Time-resolved firing activity of cold– and warm-sensitive sensory units in response to a cooling stimulus in mice (left) and naked mole-rats (right). Heatmaps display the firing rates of individual thermosensitive units, sorted by conduction velocity (CV) from lowest to highest. Each row represents one unit, with warmer colors indicating higher firing rates. Vertical dashed lines mark the onset of the cooling ramp, and the temperature profiles applied during the recordings are shown below each heatmap. AMTB was not applied in these experiments. Total number of units analyzed: mouse, cold-sensitive (47 units from 13 animals); naked mole-rat, cold-sensitive (59 units from 11 animals). **(D)** Cold-evoked firing activity in representative sensory fibers from mouse (top) and naked mole-rat (bottom) before and after AMTB (10 μM) treatment, including a final cold stimulus to assess response recovery. Traces correspond to Unit 12 (mouse) and Unit 29 (naked mole-rat), as shown in the raster plots in Figure 6H and panel B. **(E)** Raster plots of spike timing from individual cold-sensitive fibers recorded in mice before and after AMTB application. Units are grouped by conduction velocity (CV > 1.5Lm/s or < 1.5Lm/s), consistent with grouping in Fig. 6H. The trace shown in panel A corresponds to Unit 12. A total of 43 cold-sensitive units were recorded from 9 mice. **(F)** Histogram showing the distribution of conduction velocities among cold-sensitive fibers from mice (*n* = 43 units from 9 animals) and naked mole-rats (*n* = 44 units from 8 animals). **(G)** Histogram showing the distribution of soma cross-sectional areas of TRPM8⁺ neurons in dorsal root ganglia from mice and naked mole-rats. A total of 171 neurons from 3 C57BL/6 mice and 447 neurons from 3 naked mole-rats were analyzed.

